# Bleeding in cardiac patients prescribed antithrombotic drugs: Electronic health record phenotyping algorithms, incidence, trends and prognosis

**DOI:** 10.1101/538249

**Authors:** Laura Pasea, Sheng-Chia Chung, Mar Pujades-Rodriguez, Anoop D. Shah, Samantha Alvarez-Madrazo, Victoria Allan, James T. Teo, Daniel Bean, Reecha Sofat, Richard Dobson, Amitava Banerjee, Riyaz S. Patel, Adam Timmis, Spiros Denaxas, Harry Hemingway

## Abstract

**Background:** Clinical guidelines and public health authorities lack recommendations on scalable approaches to defining and monitoring the occurrence and severity of bleeding in populations prescribed antithrombotic therapy. We aimed to develop electronic health record algorithms for different bleeding phenotypes, and to determine the incidence, time trends and prognosis of bleeding in patients with incident cardiac disorders indicated for antiplatelet and/or vitamin K antagonist (VKA) therapy.

**Methods:** We examined linked primary care, hospital admission and death registry electronic health records (CALIBER 1998-2010, England) of patients with newly diagnosed atrial fibrillation, acute myocardial infarction, unstable angina or stable angina to develop algorithms for bleeding events. Kaplan-Meier plots were used to estimate the incidence of bleeding events and we used Cox regression models to assess prognosis for all-cause mortality, atherothrombotic events and further bleeding following bleeding events.

**Results:** We present electronic health record phenotyping algorithms for bleeding based on bleeding diagnosis in primary or hospital care, symptoms, transfusion, surgical procedures, and haemoglobin values. In validation of the phenotype we estimated a positive predictive value of 0.88 (95% CI: 0.64, 0.99) for hospitalised bleeding. Amongst 128,815 patients, 27259 (21.2%) had at least one bleeding event, with 5 year risks of bleeding of 29.1%, 21.9%, 25.3% and 23.4% following diagnoses of atrial fibrillation, acute myocardial infarction, unstable angina and stable angina respectively. Rates of hospitalised bleeding per 1000 patients more than doubled from 1.02 (95% CI: 0.83, 1.22) in January 1998 to 2.68 (95% CI: 2.49, 2.88) in December 2009 coinciding with increased rates of antiplatelet and VKA prescribing. Patients with hospitalised bleeding and primary care bleeding, with or without markers of severity, were at increased risk of all-cause mortality and atherothrombotic events compared to those with no bleeding. For example the hazard ratio for all-cause mortality was 1.98 (95% CI: 1.86, 2.11) for primary care bleeding with markers of severity, and 1.99 (95% CI: 1.92, 2.05) for hospitalised bleeding without markers of severity, compared to patients with no bleeding.

**Conclusions:** Electronic health record bleeding phenotyping algorithms offer a scalable approach to monitoring bleeding in the population. Incidence of bleeding has doubled in incidence since 1998, affects 1 in 4 cardiac patients, and is associated with poor prognosis. Efforts are required to tackle this iatrogenic epidemic.

What is already known?
- Clinical guidelines and public health authorities lack recommendations on how to define or monitor the occurrence and severity of bleeding in populations.
- This is particularly important because clinical guidelines increasingly recommend the use of one, two or three antiplatelet and vitamin K antagonist drugs to lower the risk of subsequent atherothrombotic events in common heart diseases including atrial fibrillation, acute coronary syndromes and chronic stable angina.
- Clinical guidelines lack consistent recommendations of how to reduce the main side effect of bleeding.
- For acute myocardial infarction it has been shown that combining primary care electronic health records (which include information from hospital discharge summaries) and hospital admission data can generate valid EHR disease phenotypes and provide real-world estimates of disease occurrence.

What is not known?
- It is not known how to define bleeding occurrence and severity in large scale, unselected populations by combining available information on bleeding diagnosis in primary or hospital care, symptoms, transfusion, surgical procedures, and haemoglobin values.
- The population-based incidence, time trends and long-term prognosis of bleeding have not been evaluated in people with common cardiac disorders.
- Comparisons of the population burden of bleeding across common cardiac disorders, such as atrial fibrillation, acute coronary syndromes and stable angina, are lacking.

What this study adds?
- **Phenotype**: We developed standardised replicable EHR phenotyping algorithms defining bleeding and severity measures based on available clinical information across structured primary and hospital care EHR sources.
- **Incidence**: At 5 years of follow-up, one in five patients with cardiac disease had a bleeding event and 6.5% had fatal or severe bleeding.
- **Trends**: There was approximately a two-fold increase in incidence of primary care and hospitalised bleeding between 1998 and 2010. The rate of fatal bleeding remained stable.
- **Prognosis**: Patients with bleeding recorded in primary care or in hospital admissions are at increased bleeding between 1998 and 2010. The rate of fatal bleeding remained stable, risk of all-cause death and atherothrombotic events.

## INTRODUCTION

Bleeding is among the most common serious side effects of modern medicine, but clinicians and health systems lack basic information on how to define and monitor the occurrence and severity of bleeding in populations. Multiple clinical guidelines make recommendations for the use of antithrombotic drugs but these guidelines have no(1, 2) recommendations about bleeding prevention, for example by concomitant proton pump inhibitor prescription.(3) Likewise public health authorities have not made recommendations for monitoring the population burden of this partially preventable cause of morbidity and mortality. There are increasing numbers of people with common heart diseases on one, two or three antithrombotic agents because of better implementation of longstanding trial evidence (e.g. aspirin in the secondary prevention of cardiovascular disease), the introduction of new drugs (e.g. P2Y12 receptor antagonists such as ticagrelor) and the prolongation (lifelong) of regimes which were initially introduced for fixed durations (e.g. prolonged use of dual antiplatelet therapy after acute myocardial infarction).(4-6)

However, the population burden (incidence, time trends and prognosis) of bleeding in people with common cardiac disorders has not been quantified. Small regional studies of intracerebral haemorrhage suggest increased incidence.(7) However, it is not clear whether the national incidence of bleeding of different severities, is increasing over time with the rise in anti-thrombotic use. Bleeding risks, often defined differently, have been described in individual diseases (atrial fibrillation(8), acute coronary syndromes(9), and stable coronary disease(10)) but there are no studies comparing risks across common cardiac disorders.

A central reason for these uncertainties is the lack of standardised definitions to measure bleeding occurrence and severity which are scalable across populations and different national health systems. We have previously shown that consistent definitions of disease and health conditions using diverse EHR can be shown to be valid and allow comparisons across countries.(11,12) Where small numbers of bleeding events are involved (e.g. in trials, or consented research cohorts(3) manual adjudication of case records is used to classify bleeding according to the BARC(13) or TIMI(14) scheme. However, such approaches are not feasible, and would be too expensive, for monitoring bleeding in large populations. Using the information obtained by linking multiple sources of electronic Health Records (EHR) across primary and hospital care may provide useful information. For example for acute myocardial infarction it has been shown that combining primary care electronic health records (which include information from hospital discharge summaries) and hospital admission data can generate valid EHR disease phenotypes and provide real-world estimates of disease occurrence.(15) Previous EHR studies of bleeding endpoints have been confined to hospitalised patients,(16-18) or one anatomical site (e.g. upper gastrointestinal bleeding(19-21)), or based on commercially insured or administrative claims data(22, 23) **(eTable 1).** However, the efficient use of information related to bleeding (e.g. diagnosis, anatomical site, fatality, length of hospital stay, haemoglobin, transfusion, endoscopy, surgical interventions) could help to generate population estimates of bleeding occurrence and severity.

We sought to address the following questions: First how can population-based EHR, spanning primary and hospital care, be used to define valid, replicable algorithms of bleeding occurrence and bleeding severity? Second, what is the long term cumulative incidence of bleeding events across patients with incident atrial fibrillation, acute myocardial infarction, unstable and stable angina who are prescribed with different antiplatelet and anticoagulation regimens? Third to what extent has the incidence of bleeding increased over time with the changes in antithrombotic management? Fourth, to what extent is bleeding of differing severity associated with long term prognosis in terms of all-cause mortality, atherothrombotic events and recurrent bleeding?

We used the CALIBER(24) research platform of linked primary, hospital, myocardial ischaemia registry and mortality data. EHR phenotypes have been developed in CALIBER for acute MI,(15) atrial fibrillation,(25) and stable coronary disease(26). Cohort studies of their associations with blood pressure,(27) diabetes,(28) smoking,(29) socioeconomic deprivation,(30) rheumatoid arthritis,(31) alcohol consumption(32) and neutrophil counts(33) have supported their validity.

## METHODS

### Linked electronic health records

We used data from the CALIBER(24) resource. CALIBER links EHR from primary care general practices (Clinical Practice Research Datalink [CPRD]), hospital admissions (Hospital Episode Statistics [HES]), myocardial ischaemia registry (Myocardial Ischaemia National Audit Project [MINAP]) and cause-specific mortality (Office for National Statistics [ONS]) data in England. The 4% sample of the England’s population in CPRD available for linkage is representative in terms of age, sex and overall mortality.(34-36) In CALIBER, EHR disease phenotypes(37)have been developed through collaborations between clinicians, epidemiologists and statisticians and a number of risk factors and cardiovascular and non-cardiovascular disease endpoints have been validated for cardiovascular research.(15, 24-33) The study was approved by the Independent Scientific Advisory Committee of the Medicines and Health care products Regulatory Agency in the UK, protocol number 14_133.

### Study population

The study population consisted of patients with cardiac disease, i.e. those who were potential candidates for antiplatelet and/ or vitamin K antagonist (VKA) therapy, in CALIBER during 1997-2010. To define this population we used pre-existing validated disease phenotypes **[https://www.caliberresearch.org/portal]**. Patients were eligible if they were aged 18 years and above and entered the cohort at their first diagnosis of atrial fibrillation, acute myocardial infarction, unstable angina or stable angina in primary or hospital care records. They were followed up until death, transfer out of their primary care practice (i.e. loss to follow-up), or the date of administrative censoring (March 2010).

We analysed baseline characteristics of patients stratified by initial cardiac disease. Using prescribing data we summarised therapy duration (median and interquartile range days) of between cohort entry and first bleeding event. To calculate duration, a patient’s prescription was assumed to be continuous if issued within 90 days of the previous one (90 days is the longest allowed duration of prescriptions in the UK). Treatments were grouped as aspirin monotherapy, adenosine diphosphate (ADP) receptor inhibitor monotherapy, dual antiplatelet therapy (aspirin and ADP receptor inhibitor), VKA monotherapy, VKA and one antiplatelet (aspirin <u>or</u> ADP receptor inhibitor) and triple therapy (VKA, aspirin <u>and</u> ADP receptor inhibitor).

### Electronic health record data relevant to definition of bleeding phenotypes

Within CALIBER, bleeding events were captured in primary care data (Read terms), hospital admissions administrative data (ICD-10 terms) and death registry (ICD-9 and ICD-10 terms) **(eTable 2).** The description of the terms used contained information on anatomical site of bleeding. Hospital records indicated the diagnosis position (i.e. primary or secondary reason for hospitalisation) and the length of hospitalisation was calculated using admission and discharge dates. Procedures relevant to bleeding (transfusion, bleeding surgical interventions and endoscopy) were captured in hospitalisation records using OPCS codes. Drug prescriptions were available in primary care data, classified according to British National Formulary (BNF) chapter. Clinical biomarkers such as haemoglobin, were also captured in primary care.

### Algorithmic combinations to define bleeding EHR phenotypes

The construction of the CALIBER bleeding EHR phenotype **(Figure 1)** is fully explained in **eMethods.** In short, we applied a structured approach to phenotyping, previously demonstrated by Morley et al,(25) involving iterative steps of diagnosis code reviews, descriptive analyses, and expert input. We used published trial protocols of definition of major bleeding(13,14, 38) to identify candidate markers of bleeding severity. We included the sub-set of markers which were available in the EHR (for example the HES data does not record haemoglobin measurements), and evaluated associations with short-term mortality in order to develop the severe bleeding EHR phenotype. We defined **fatal bleeding** as a bleeding cause of death (underlying or otherwise) in the national death registry or all-cause death within 7 days of a bleeding record in primary or hospital care. We identified 4 markers of bleeding severity available within our data: 1) bleeding as a primary reason for hospitalisation combined with at least 14 days hospitalisation; 2) bleeding site (intracranial, ruptured aortic aneurysm or haemopericardium; 3) bleeding from more than one site on the same day; and 4) a transfusion record in hospital care within 30 days of a bleeding record.

**Figure 1:**
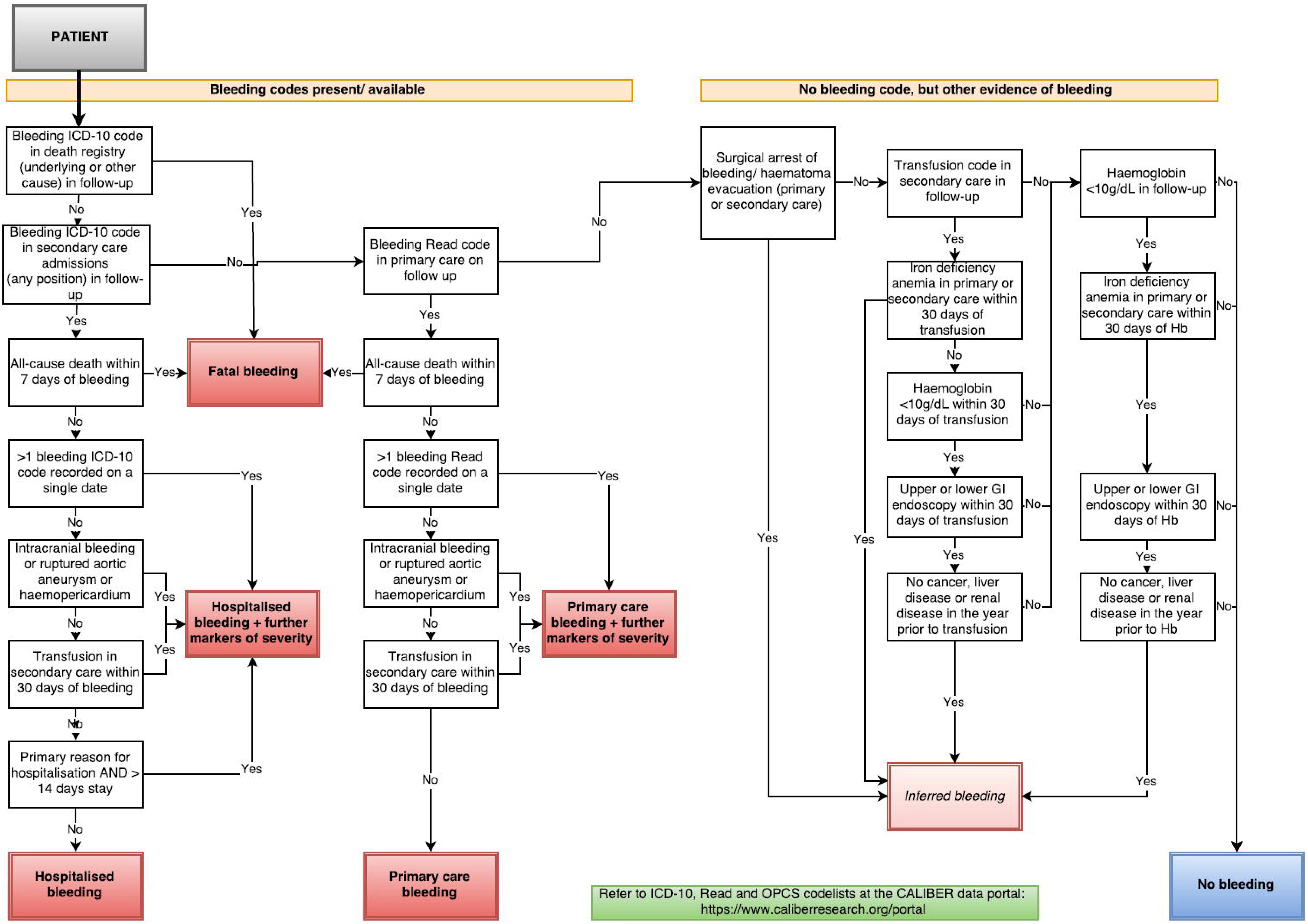
Bleeding EHR phenotype algorithm for fatal, hospitalised, primary care and inferred bleeding with and without additional markers of severity

We classified non-fatal bleeding events as hospitalised or primary care with further markers of severity (henceforth referred to as **‘hospitalised+MS’** and **‘primary care+MS’),** and hospitalised or primary care without markers of severity (referred to as **‘hospitalised’** and **‘primary care’).** For patients with no bleeding code in either primary care or hospital records, possible bleeding events may be inferred where there are records that provide evidence suggesting bleeding, for example transfusions and low haemoglobin.

We validated the phenotype algorithm through manual case note review. Two clinicians (blinded to the ICD-10 and OPCS-4 codes recorded) reviewed the entire hospital record (charts, referral letters, discharge letters, imaging reports) for 283 patient hospital episodes from two large NHS Trusts (University College London Hospitals NHS Foundation Trust and Kings College Hospital NHS Foundation Trust). The hospital record corpus (14,364,947 words in total) was made available as a single text files per patient, through the use of CogStack(39), method of enterprise-wide retrieval and extraction architecture for structured and unstructured information which integrates data across multiple EHR systems in a hospital. Patient consent for reviewing these records was provided from the NIHR funded SIGNUM study of stroke patients. Bleeding assignments from the clinicians’ review was compared with those from the bleeding algorithm and we estimated the positive predictive value (PPV), negative predictive value (NPV), sensitivity and specificity using the case review data as the gold standard.

### Statistical analysis

#### Cumulative bleeding incidence in four cardiac diseases

The incidence of any bleeding and fatal, hospitalised+MS or primary care+MS bleeding was assessed using Kaplan-Meier plots stratified by cardiac disease type (atrial fibrillation, acute myocardial infarction, unstable angina or stable angina).

#### The association between antithrombotic prescribing and bleeding

Cox proportional hazard models were used to estimate hazard ratios for the association between antithrombotic therapies and first bleeding event of any severity and fatal or bleeding+MS event. Antithrombotic therapy prescriptions were included in the models as a time-dependent variable. Possible states were no antithrombotic therapy (the reference group), aspirin, ADP receptor inhibitor, dual antiplatelet therapy, vitamin K antagonist, vitamin K antagonist and one antiplatelet (aspirin or ADP receptor inhibitor), and triple therapy. Patients were followed up until their first bleeding event of any severity and until their first fatal or bleeding+MS event. The Cox models were adjusted for age and sex.

#### Time trends in bleeding

We estimated the number of fatal, hospitalised+MS, primary care+MS, hospitalised and primary care bleeding events per 1000 patients at monthly intervals between 1997 and 2010. To do this we divided the number of bleeding events recorded by the total number of patients at risk each month. Loess smoothed lines were fitted to detect changes in incidence over time. Similarly, we estimated the time trends for the number of antithrombotic prescriptions issued each month.

#### Prognosis following bleeding

We used Cox proportional hazard models to estimate hazard ratios (HR) for the association between first bleeding events, all-cause mortality, and atherothrombotic events (composite of cardiovascular death, ischaemic or unspecified stroke, or myocardial infarction). Bleeding severity (hospitalised+MS, primary care+MS, hospitalised, primary care and inferred) was treated as a time-dependent variable in the models to prevent immortal time bias. The possible bleeding variable states were no bleeding (reference group), primary care, primary care+MS, hospitalised or hospitalised+MS. All patients started follow up in the no bleeding state and changed to the relevant bleeding state at the time of their first bleeding event. Models were also adjusted for age, sex and baseline disease history (diabetes, stroke, peripheral arterial disease, cancer, renal disease, peptic ulcer, bleeding diatheses, chronic anaemia). We also explored the risk of recurrent bleeding in the subgroup of patients that had non-fatal bleeding events using Kaplan-Meier plots, following patients from the time of their first non-fatal bleeding event.

#### Modelling assumptions

The proportional hazards assumptions of Cox models were checked using residual and log(−log) plots.

All analyses were performed using R version 3.2.

### Patient involvement

No patients were involved in setting the research question and study outcome or the design and implementation of the study. There are no current plans to disseminate results with patient groups.

## RESULTS

### Study population

Our study population consisted of 128,815 patients in 224 general practices newly diagnosed with atrial fibrillation, acute myocardial infarction, unstable angina and/or stable angina between 1997 and 2010. They were followed up for a total of 559,161 person-years, a median of 3.7 years (IQR: 1.5, 6.9). The mean age was 71.5 years at cohort entry (43.8% aged ≥75 years) and 48.5% were women.

Patient characteristics stratified by cardiac disease are shown in **Table 1.** Atrial fibrillation patients were older than the coronary disease patients and the majority were women. In contrast the coronary disease patients were mostly men. The atrial fibrillation patients also had higher prevalence of history of stroke, renal disease, cancer and chronic anaemia. The majority of patients in all four disease groups were prescribed at least 1 antithrombotic drug between cohort entry and first bleeding event or end of follow up in those who did not bleed.

**Table 1:** Baseline characteristics of people with four common cardiac diseases

|  | **Atrial Fibrillation**<br>**(n= 27,061)** | **Myocardial**<br>**Infarction**<br>**(n=25,031)** | **Unstable Angina**<br>**(n=9,500)** | **Stable Angina**<br>**(n= 67,223)** |
| --- | --- | --- | --- | --- |
| **Demographics** |  |  |  |  |
| Age (years), mean (SD) | 76.6 (12.8) | 69.9 (13.5) | 69.1 (13.2) | 70.4 (12.3) |
| Age≥75years, n(%) | 16946 (62.6) | 9982 (39.9) | 3488 (36.7) | 26059 (38.8) |
| Women, n (%) | 14266 (52.7) | 9206 (36.8) | 4169 (43.9) | 31365 (46.7) |
| Highest quintile of deprivation<br>(most deprived), n (%) | 5137 (19.0) | 4758 (19.1) | 1947 (20.5) | 13837 (20.6) |
| **Behaviours** |  |  |  |  |
| Current smoker, n (%) | 2277 (10.5) | 3691 (19.5) | 1058 (14.0) | 6229 (11.4) |
| History of alcohol abuse, n (%) | 2627 (9.7) | 2430 (9.7) | 908 (9.6) | 6459 (9.6) |
| **Medical history prior to cohort**<br>**entry <sup>a</sup>** |  |  |  |  |
| Type 2 Diabetes, n (%) | 2695 (10.0) | 2922 (11.7) | 1222 (12.9) | 8728 (13.0) |
| Ischaemic or unspecified stroke,<br>n (%) | 2169 (8.0) | 1462 (5.8) | 558 (5.9) | 3435 (5.1) |
| Peripheral arterial disease, n (%) | 2276 (8.4) | 2147 (8.6) | 865 (9.1) | 6199 (9.2) |
| Renal disease, n (%) | 2570 (9.5) | 1731 (6.9) | 694 (7.3) | 4351 (6.5) |
| Non-metastatic cancer, n (%) | 5427 (20.1) | 3158 (12.6) | 1155 (12.2) | 8701 (12.9) |
| Metastatic cancer, n (%) | 526 (1.9) | 209 (0.8) | 74 (0.8) | 520 (0.8) |
| Peptic ulcer, n (%) | 1814 (6.7) | 1713 (6.8) | 753 (7.9) | 5074 (7.5) |
| Bleeding diatheses and<br>coagulation disorders, n (%) | 312 (1.2) | 175 (0.7) | 77 (0.8) | 534 (0.8) |
| Chronic anaemia, n (%) | 4982 (18.4) | 2808 (11.2) | 1198 (12.6) | 8125 (12.1) |
| **Biomarkers at cohort entry <sup>b</sup>** |  |  |  |  |
| SBP (mmHg), mean (SD) | 140 (21.8) | 143 (21.2) | 142 (21.2) | 142 (20.5) |
| % Missing | 29.7 | 33.2 | 25.8 | 21.6 |
| Haemoglobin (g/dL), mean (SD) | 12.9 (1.97) | 13.5 (1.91) | 13.5 (1.75) | 13.6 (1.69) |
| % Missing | 56.2 | 65.9 | 62 | 59.1 |
| Creatinine (mmol/l), mean (SD) | 107 (59.1) | 108 (56.1) | 105 (55.9) | 102 (46.1) |
| % Missing | 49.5 | 57.6 | 54.1 | 49.7 |
| **Antithrombotic therapies (n, %) and duration (median, IQR) during follow-up <sup>c</sup>** |  |  |  |  |
| Any antithrombotic therapy | 16868 (62.3) | 19950 (79.7) | 7947 (83.7) | 55619 (82.7) |
| Aspirin monotherapy | 10787 (39.9) | 16511 (66.0) | 6695 (70.5) | 48262 (71.8) |
| Duration (days) | 382 (114, 908) | 791 (267, 1742) | 765 (268, 1691) | 842 (305, 1752) |
| ADP receptor inhibitor monotherapy | 1264 (4.7) | 3683 (14.7) | 1425 (15.0) | 7351 (10.9) |
| Duration (days) | 150 (46, 486) | 94 (30, 376) | 121 (42, 495) | 181 (52, 652) |
| Dual antiplatelet therapy | 1594 (5.9) | 8673 (34.6) | 2417 (25.4) | 9539 (14.2) |
| Duration (days) | 186 (90, 426) | 349 (143, 478) | 272 (98, 488) | 261 (90, 476) |
| VKA monotherapy | 7149 (26.4) | 1666 (6.7) | 853 (9.0) | 6287 (9.4) |
| Duration (days) | 427 (146, 1083) | 216 (82, 626) | 318 (110, 844) | 344 (113, 938) |
| VKA + 1 antiplatelet | 3003 (11.1) | 1426 (5.7) | 637 (6.7) | 3892 (5.8) |
| Duration (days) | 85 (51, 163) | 106 (55, 262) | 90 (54, 228) | 90 (51, 214) |
| VKA + 2 antiplatelets | 266 (1.0) | 321 (1.3) | 102 (1.1) | 430 (0.6) |
| Duration (days) | 68.5 (39, 93.2) | 68.0 (43, 116.0) | 62.5 (39, 90.0) | 57.0 (35, 84.0) |

**Note:** SD=standard deviation, SBP=systolic blood pressure, BMI= body mass index, IQR= interquartile range, ADP=adenosine diphosphate, VKA=vitamin K antagonist
<sup>a</sup> Any record prior to cohort entry
<sup>b</sup> Nearest record to entry within 1 year prior to entry
<sup>c</sup> Between cohort entry and 1<sup>st</sup> bleeding event or end of follow-u

### Applying the CALIBER bleeding EHR phenotype algorithm

The bleeding algorithm is shown in **Figure 1.** We identified 39,804 bleeding records from 27,259 (21.2%) patients in our cohort. 59.4% of coded bleeding events were captured in primary care, 50.2% in hospital admissions, and 3.8% events in the death registry. Allowing a 30 day window, only 13.2% of coded bleeding events were captured in 2 or more data sources. The overlap of bleeding events between the data sources used is shown in **eFigure 1.** For hospitalised bleeding we estimated a PPV of 0.88 (95% CI: 0.64, 0.99), a NPV of 0.94 (0.90, 0.97), a sensitivity of 0.48 (0.30, 0.67) and a specificity of 0.99 (0.97, 1.00) **(eTable 3).**

We identified 1,492 further possible bleeding events occurring in 1,144 patients with no bleeding diagnosis recorded in primary care or hospital records through the following routes: (1) transfusion and presence of iron deficiency anaemia diagnosis within 30 days (n=689); (2) surgical procedures to arrest bleeding or for haematoma evacuation (n=477); (3) haemoglobin<10g/dL, iron deficiency anaemia diagnosis and endoscopic examination within 30 days and no cancer, liver or renal disease records in the previous year (n=249); (4) transfusion, haemoglobin<10g/dL and endoscopic examination within 30 days and no cancer, liver or renal disease records in the year prior (n=77).

### Cumulative incidence of any bleeding and fatal bleeding or bleeding with markers of severity

At 5 years 29.1% (95% CI: 28.2, 29.9%) of atrial fibrillation patients, 21.9% (21.2, 22.5%) of myocardial infarction patients, 25.3% (24.2, 26.3%) of unstable angina patients and 23.4% (23.0, 23.8%) of stable angina had bleeding of any kind **(Figure 2).** Risks of fatal bleeding, hospitalised+MS or primary care+MS bleeding events at 5 years were 9.9% (9.3, 10.4%) for atrial fibrillation patients, 6.1% (5.8, 6.5%) for myocardial infarction patients, 6.8% (6.0, 7.2%) for unstable angina patients and 5.7% (5.5, 5.9%) for stable angina.

**Figure 2:**
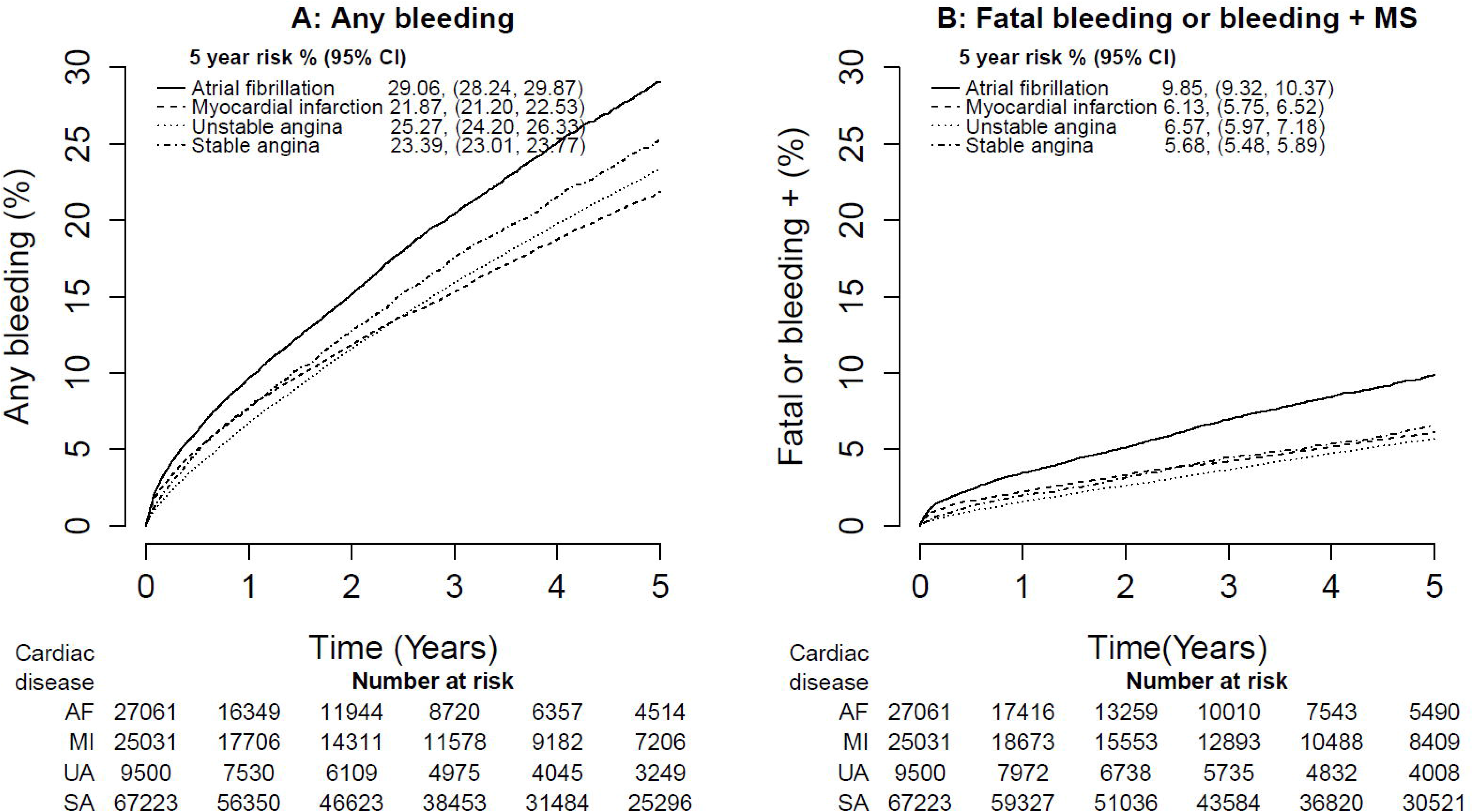
Five year risk of CALIBER bleeding from time of initial atrial fibrillation, acute myocardial infarction, unstable angina or stable angina (n= 128,815 patients); A: any bleeding, B: Fatal bleeding or bleeding with further markers of severity

**Note:** MS= markers of severity; In panel A ‘Any bleeding’ includes fatal, hospitalised+MS, hospitalised, primary care+MS and primary care bleeding events; in panel B ‘Fatal bleeding or bleeding +MS’ includes fatal, hospitalised+MS and primary care+MS bleeding events only

### Time trends in bleeding incidence and antithrombotic prescribing

The estimated number of hospitalised+MS bleeding events per 1000 active patients increased from 0.32 (0.24, 0. 40) in January 1998 to 0.54 (0.45, 0.62) in December 2009. Contrarily, in primary care+MS bleeding events per 1000 active patients decreased from 0.80 (95% CI: 0.70, 0.91) in January 1998 to 0.34 (0.23, 0.45) in December 2009. The incidence of fatal bleeding remained steady **(Figure 3A).**

**Figure 3:**
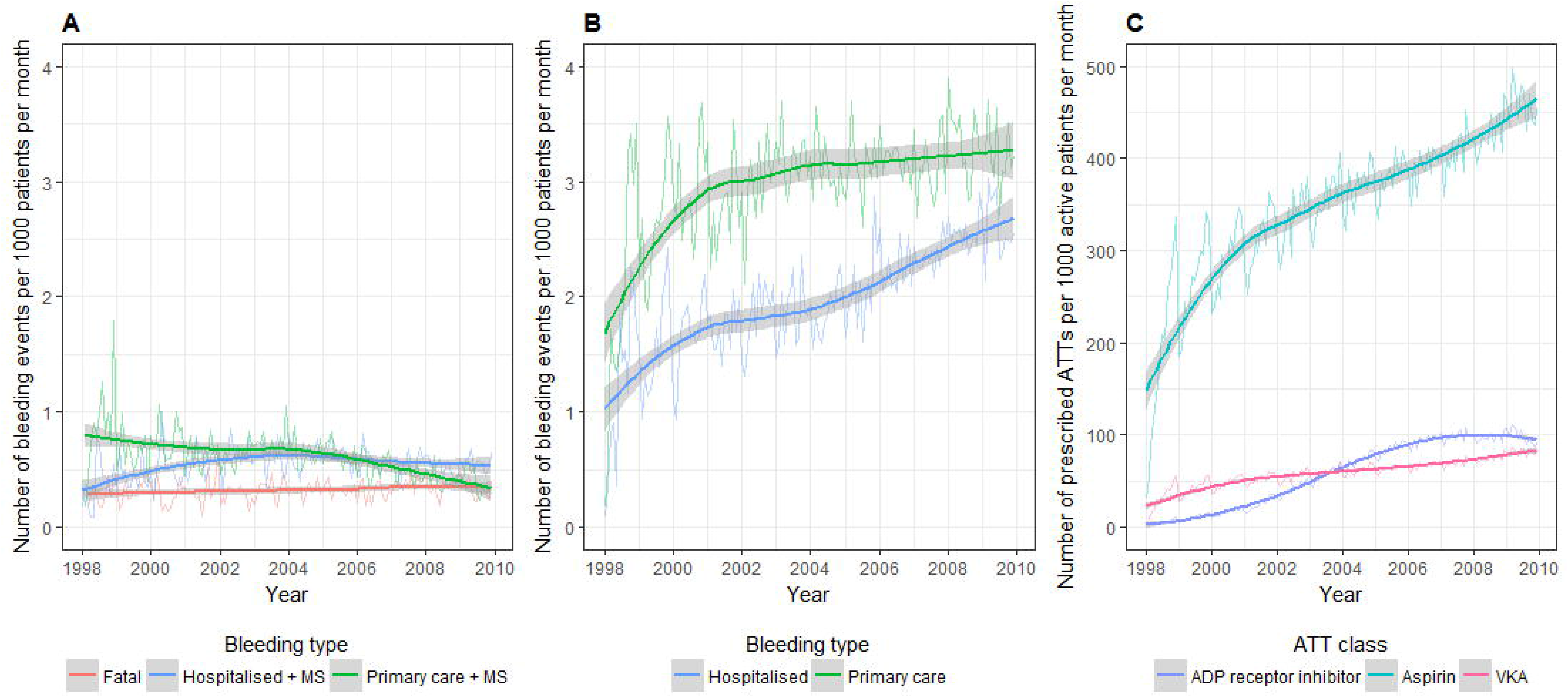
Time trends of fatal, hospitalised and primary care bleeding events and antithrombotic prescribing 1998-2010 in CALIBER. A: Fatal, hospitalised +MS and primary care +MS bleeding events; B: Hospitalised and primary care bleeding events; C: Prescriptions for ADP receptor inhibitors, aspirin and vitamin K antagonists.

**Note:** Fitted lines are loess smoothed curves with shaded 95% confidence intervals; MS= markers of severity; ATT = antithrombotic therapy; VKA= vitamin K antagonists

There were increases in hospitalised and primary care bleeding events without markers of severity **(Figure 3B).** The estimated number of hospitalised bleeding events per 1000 active patients increased from 1.02 (0.83, 1. 22) in January 1998 to 2.68 (2.49, 2.88) in December 2009, and for primary care bleeding events the increase was from 1.70 (1.44, 1.95) to 3.31 (3.06, 3.57). This corresponded to the rise of rates of prescribed antithrombotic therapies over the study period **(Figure 3C).** From January 1998 to December 2009, the increase in the number of prescriptions issued per 1000 active patients for aspirin, ADP receptor inhibitor, and VKA was 147.9 (95% CI: 127.4, 168.3) to 465.1 (444.6, 485.6), 2.8 (0.2, 5.4) to 94.8 (92.2, 97.4), 22.7 (19.2, 26.1) to 83.7 (80.2, 87.1), respectively.

Overall, patients prescribed with more aggressive antithrombotic therapies (dual antiplatelet therapy, vitamin K antagonists, and triple therapy) had significantly higher risk of bleeding events compared with those not prescribed antithrombotic therapies **(Figure 4).** Compared with those not prescribed antithrombotic therapies, patients who were prescribed triple therapy had 3.4 (2.6, 4.4) times increased risk of any bleeding and 5.7 (3.7, 8.7) times increased risk of fatal or bleeding+MS events.

**Figure 4:**
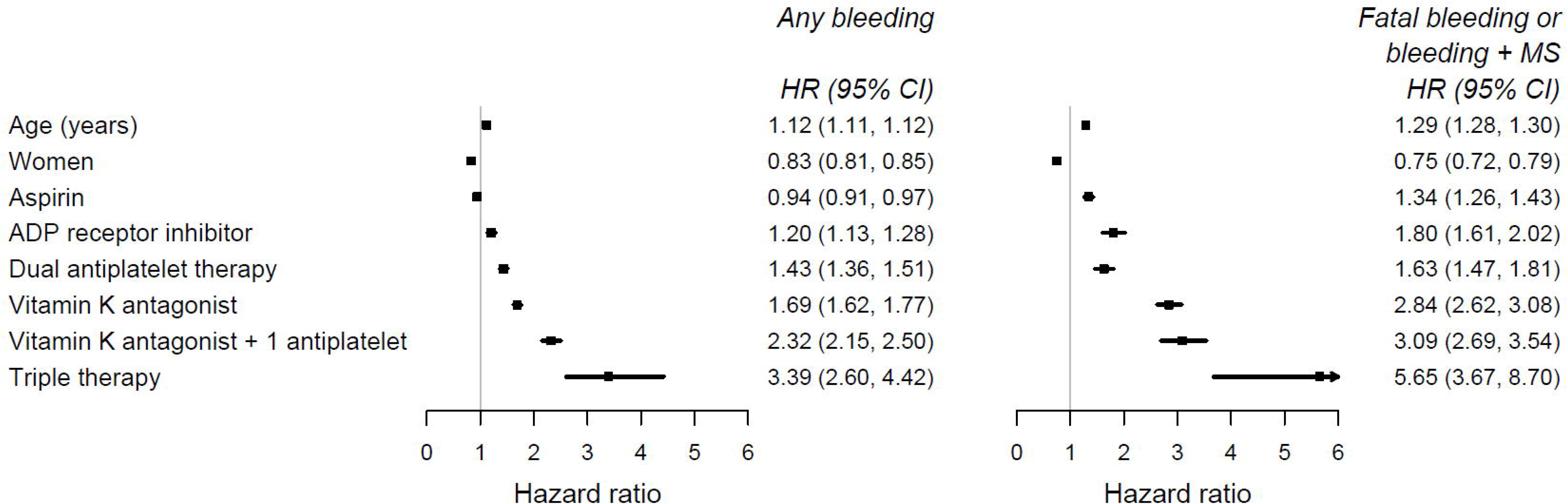
The association between antithrombotic therapy prescribing and any bleeding and fatal or bleeding + MS events adjusted for age and sex

**Note:** HR= Hazard ratio; MS= markers of severity;

### Death, atherothrombotic events following first bleeding event

Patients were at increased risk of all-cause mortality and cardiovascular death, stroke or myocardial infarction following their first bleeding event and this association was observed across all bleeding severities **(Figure 5).** Based on the magnitude of relative risks for prognostic outcomes three levels of bleeding severity were identified: class I: The greatest prognostic risk was observed in hospitalised+MS bleeding (class I), followed by hospitalised or primary care+MS, or Inferred bleeding (class II). The lowest prognostic risk was associated with primary care bleeding (class III).

**Figure 5:**
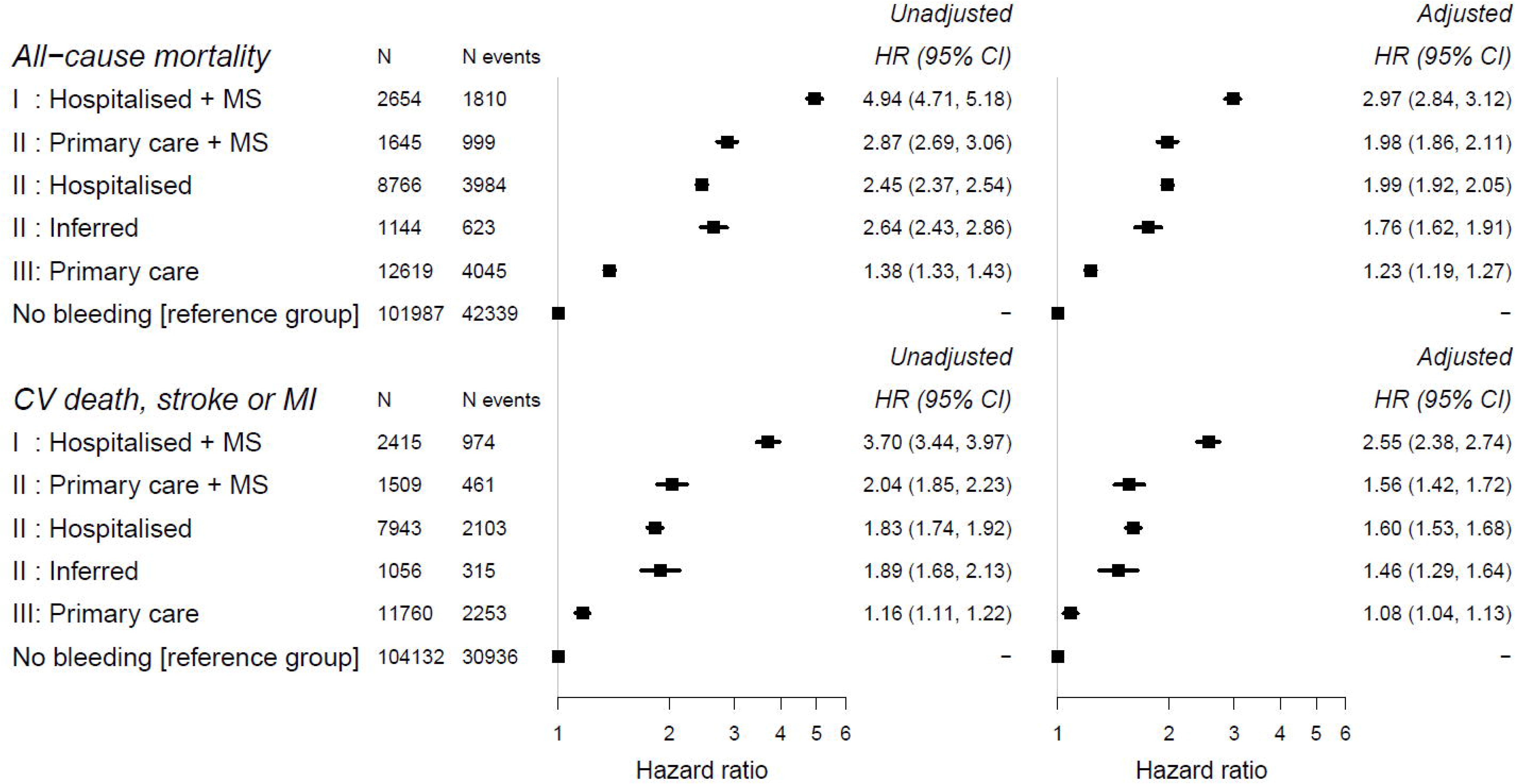
The association between non-fatal bleeding severity classes and all-cause mortality and cardiovascular death, stroke or myocardial infarction (vs. no bleeding)

**Note:** Adjusted estimates are adjusted for age, sex and comorbidities; MS= markers of severity; HR= hazard ratio; CI= confidence interval; CV= cardiovascular; Ml= myocardial infarction

Compared to patients with no bleeding, the adjusted HR for all-cause mortality was 2.97 (2.84, 3.12) for class I bleeding and 1.23 (1.19, 1.27) for class III bleeding. Similarly, the adjusted HR for cardiovascular death, stroke or MI events was 2.55 (2.38, 2.74) for class I and 1.08, (1.04, 1.13) for class III bleeding.

Risk of recurrent bleeding increased following an initial bleeding event **(eFigure 2).** The cumulative risks were greater if the initial bleeding event had further markers of severity. The five-year recurrent event rates of any bleeding and fatal, hospitalised+MS or primary care+MS bleeding were 32.4% (31.8, 33.0), and 8.3% (7.9, 8.6) respectively. Among patients who initially experienced a bleeding event with markers of severity, their 5 year recurrent event rate was 37.4% (36.0, 38.8) for any bleeding and 23.1% (21.9, 24.3) for fatal, hospitalised+MS or primary care+MS bleeding.

## DISCUSSION

In a population based study of linked primary care and hospital EHR in 128,815 patients with newly diagnosed common cardiac disorders we found that bleeding has doubled in incidence since 1998, affects 1 in 4 cardiac patients, and is associated with poor prognosis in terms of all-cause mortality and subsequent atherothrombotic events. This suggests that bleeding has become an epidemic, most likely iatrogenic, caused by the increase in the use of antithrombotic medication. The phenotype algorithms made available here distinguish three prognostic classes of bleeding severity which may be used by health systems and public health authorities to focus efforts to tackle the growing population impact of bleeding on health outcomes.

### Bleeding EHR phenotype algorithm: importance of linked electronic health records

We developed standardised and replicable EHR phenotyping algorithms for bleeding and severity measures based on available clinical information across primary and hospital care. The algorithms combine information on diagnoses, procedures, transfusion and haemoglobin. Unlike previous EHR studies which defined bleeding events using bleeding codes only, we demonstrated the depth of information readily available within linked EHR and the capability to achieve a more granular case definition by combining diagnoses terms with continuous measurements. Our results highlighted the importance of using multiple linked data sources for defining and validating the bleeding phenotype in EHR. No individual data source used in this study had complete coverage of coded bleeding diagnoses, transfusions, causes of death and other bleeding relevant data, and only 13.2% of bleeding cases were captured in multiple data sources **(eFigure 1).** Individual components of the phenotype, such as subgroups of the bleeding codes, have been validated in previous studies in CPRD,(21) HES(20) and other electronic health record systems,(16-19, 22,23) and our analysis of outcomes following bleeding adequately reflected expected results across levels of bleeding severity. It has been previously shown that using hospital discharge coding alone misses bleeding events compared with a manual review of case notes;(3) nonetheless our use of multiple sources of EHR led to estimation of a higher incidence of bleeding at one year than in the study with manual case note review.

### Validation of bleeding phenotype

We provide new evidence of the validity of ICD-10 codes used in our bleeding EHR phenotype algorithm. We found a PPV of 0.88, i.e. 88% of bleeding events identified by these codes were indeed bleeding events according to the independent review of the entire hospital record by two clinicians, blinded to the ICD-10 code assignment. We also provide evidence that these hospital codes underestimate the occurrence of bleeding, with a sensitivity estimate of 0.48, consistent with Li et al(3) who found that of 38% of hospitalised bleeding events in a cohort of patients with vascular disease were not captured using clinical codes. This suggests that our study might have underestimated the incidence of bleeding amongst patients prescribed antithrombotic therapies. There have been few previous studies of the validity of ICD-10 codes in the UK against full review of hospital records, partly due to the difficulties in accessing the hospital records; our informatics approach using CogStack (39) for validation is scalable, replicable, rapid and low cost.

### Bleeding EHR phenotype: inferring bleeding events

A previous study showed that it is appropriate to infer disease cases in EHR where diagnosis codes are absent.(25) We identified 1,144 patients with no coded bleeding diagnosis present but exhibiting signs or symptoms of bleeding, such as low haemoglobin, iron-deficiency anaemia or with a recorded bleeding related procedure, excluding cases where bleeding may not be the cause of these signs, symptoms and procedures (i.e. cancer, liver and renal diseases). This highlights the potential of looking beyond diagnosis codes in EHR to obtain more accurate estimates of bleeding in safety studies of antithrombotic use. This method requires validation and cases identified using this method should be considered possible bleeding events and not definite.

### Bleeding incidence in cardiac disease populations

At 5 years of follow-up one in four patients with cardiac disease had any bleeding event and 6.5% had fatal or severe bleeding. We provided a direct comparison of bleeding within 4 cardiac diseases with varying degrees of antithrombotic use **(eTable 4).** Atrial fibrillation had highest bleeding 5-year rates both for any bleeding (29.1%) and fatal, hospitalised+MS or primary care+MS bleeding (9.9%). This is likely to reflect the higher use and longer duration of prescribed VKA and dual, triple therapy in atrial fibrillation patients. However the incidence of bleeding in myocardial infarction, unstable angina and stable angina patients was still relatively high.

### Time trends in bleeding rates over the study period

So far as we are aware there have been no previous studies evaluating the time trends in bleeding incidence in common cardiac disorders. In our study, we found that the rates of hospitalised bleeding per 1000 patients more than doubled from 1.02 in 1998 to 2.68 in 2009. We hypothesised that the increased use of antithrombotic therapies during this period would be associated with an increased incidence of bleeding. We indeed did identify increases in rates of hospitalised+MS, hospitalised and primary care bleeding events over time, consistent with an increase over the same time period. However, based on the results of our study, we cannot distinguish the relative contributions to the observed increase in bleeding incidence of the increasing range of available antithrombotic therapies, widening indications and changing guidelines for their use over time. Because hospitals receive reimbursement based on the ICD codes at discharge (40)it is possible that the observed increase in rate of bleeding is partly artefactual, i.e. due to better recording over time. However, there are 3 lines of evidence against such an artefact: 1) we also observed increases in the rate of bleeding in an entirely separate source of primary care data (reimbursed for coding relating to NICE Quality and Outcomes Framework indicators, in which bleeding is not included) used in real time clinical decision making. 2) this increase is consistent with previous evidence, of the increase in rates of intracerebral haemorrhage in the UK between 1981 and 2006.(7) and 3) prescribing of anti-thrombotic therapies, which is known to increase the risk of bleeding complications, has increased during the study period.

### Prognosis following bleeding

These bleeding events were associated with poor outcomes suggesting an increasing burden of bleeding on healthcare systems and costs in England. Our analysis of prognosis following a non-fatal bleeding event identified 3 distinct levels of severity; I: Hospitalised+MS, II: Hospitalised, Primary care +MS, or inferred bleeding and III: Primary care **(Figure 5).** This goes beyond the usual dichotomised classification of bleeding as either major or minor that is commonly reported. Increased bleeding severity was strongly associated with increased risks of all-cause mortality and atherothrombotic events. In particular we found that bleeding diagnosed in primary care, without acute hospitalisation, was associated with adverse prognosis, both as class II and as class III (with and without associated markers of severity, respectively). Thus all types of bleeding captured by the phenotype are clinically relevant. The term ‘minor bleeding’ may be misleading for clinicians, suggesting that no further action is required; while our study suggests that even a bleed in primary care without additional markers of severity is associated with 23% increased risk of death. While we have identified associations between bleeding and prognosis, in our present analyses we cannot claim these associations to be causal.

### Limitations of EHRs

EHRs have strengths and limitations for defining bleeding. Strengths include the availability of relevant, constantly updated information, at nationally representative scale, with the opportunities for international comparison(12) and the low cost of acquiring the information. The key limitation is that at national scale information is lacking on acute haemoglobin change, the number of units transfused and other details of bleeding to support the classification of bleeding severity. In clinical practice these markers are used to assess bleeding severity and have high prognostic value.(41) Their addition to EHR phenotypes, would be an important refinement to bleeding definitions. We showed some evidence that haemoglobin drop might contribute to defining bleeding severity, but our data lacked haemoglobin values measured within hospital admissions. The prescribing data reported here was confined to primary care and did not include drugs prescribed during hospitalisation or over the counter aspirin. Therefore the rates of prescribing reported may underestimate the true rates.

### Clinical implications

Our study provides evidence of an iatrogenic epidemic, demonstrating the public health burden of increasing bleeding incidence and adverse prognosis, and suggests three clinical implications. First, clinicians should ensure that the decision to prescribe antithrombotic therapy is based on a personalised evaluation of both bleeding risk and atherothrombotic risk in combination with trial results. Such an approach tailors drug treatment decisions to an individual’s expected net benefit, and is able to incorporate a patient’s utility (or disutility) from bleeding and atherothrombotic events.(42) Previous studies, for example in the setting of prolonged dual antiplatelet therapy,(42) have demonstrated the validity, and feasibility (with web calculators) of such an approach using readily available clinical data. Second, clinicians should be aware that patients who experience bleeding events, even those which are not hospitalised, are at particularly high risk and may warrant more intense monitoring. Third, we propose that bleeding events are continually monitored and reported by organisations as part of quality of care and outcome reporting not just in single cardiovascular diseases, but across whole health systems and whole populations. Indeed one general population survey of adults aged 45-75 years conducted in the US, reported antiplatelet use in 47% despite the small proportion of participants with established cardiovascular disease.(43) We have shown that the severe bleeding EHR phenotypes reported here closely match the endpoints used in trials.(26). This suggests that linked EHR can be used in ongoing reporting to estimate real-world impact of interventions.

### Future research

International standards for the EHR definition of bleeding occurrence and severity using available national and regional clinical records and based on the approach described here should be developed. Transparent reporting of EHR phenotype algorithms is required in order to make bleeding research more replicable and to compare the incidence and prognosis of bleeding of different severities in different countries and across different health systems.(12) This is important to understand the extent to which, if any, newer antithrombotic agents such as direct oral anticoagulants and ticagrelor are halting the trend of increased incidence of bleeding or reducing the severity of bleeding events. The methods validation of disease code based EHR phenotypes against the full hospital record reported here are scalable to other diseases and other hospitals.

### Conclusion

Bleeding is a major public health problem; it is common in patients with cardiac conditions, the incidence of hospitalisation for bleeding is increasing, and it is associated with high mortality. The comprehensive and reproducible bleeding EHR phenotype with three levels of severity that we have developed is informative in mortality, risk of fatal or non-fatal atherothrombotic events, and recurrent bleeding. It can be used and further developed in EHR studies of bleeding outcomes or antithrombotic safety.

## Funding

This investigators on this study were supported by multiple funding sources including, the Medical Research Council Prognosis Research Strategy (PROGRESS) Partnership (HH, grant G0902393/99558), Medical Research Council Population Health Scientist Fellowship (S-CC: grant MR/M015084/1), and by awards to establish the Farr Institute of Health Informatics Research, London and Scotland from the Medical Research Council,

Arthritis Research UK, British Heart Foundation, Cancer Research UK, Chief Scientist Office, Economic and Social Research Council, Engineering and Physical Sciences Research Council, NIHR, National Institute for Social Care and Health Research, and Wellcome Trust (LP, S-CC, MPR). LP was supported by an AstraZeneca PhD studentship. The views expressed in this paper do not necessarily represent the views of the funding bodies. LP had full access to the data and takes responsibility for the integrity of the data and the accuracy of the data analysis. All authors had final responsibility for the decision to submit for publication.

HH is a National Institute for Health Research (NIHR) Senior Investigator. His work is supported by: 1. Health Data Research UK, which is funded by the UK Medical Research Council, Engineering and Physical Sciences Research Council, Economic and Social Research Council, Department of Health and Social Care (England),

Chief Scientist Office of the Scottish Government Health and Social Care Directorates, Health and Social Care Research and Development Division (Welsh Government), Public Health Agency (Northern Ireland), British Heart Foundation and Wellcome Trust. 2. The BigData@Heart Consortium, funded by the Innovative Medicines Initiative-2 Joint Undertaking under grant agreement No. 116074. This Joint Undertaking receives support from the European Union’s Horizon 2020 research and innovation programme and EFPIA; it is chaired, by DE Grobbee and SD Anker, partnering with 20 academic and industry partners and ESC. 3. The National Institute for Health Research University College London Hospitals Biomedical Research Centre.

ADS is supported by the National Institute for Health Research University College London Hospitals Biomedical Research Centre

This study represents independent research part funded by the National Institute for Health Research (NIHR) Biomedical Research Centre at South London and Maudsley NHS Foundation Trust and King’s College London and the NIHR Biomedical Research Centre at University College London Hospitals. The views expressed are those of the author(s) and not necessarily those of the NHS, the NIHR or the Department of Health and Social Care

## Authors’ contributions

LP, SCC, MPR, AS, SAM, VA, JTT, DB, RS, RD, AB, RSP, SD, HH contributed to the idea and design of the study. LP, JTT, DB extracted and prepared the data for analysis. LP performed the analysis. LP and HH drafted the manuscript, with revisions by SCC, MPR, AS, SAM, VA, JTT, DB, RS, RD, AB, RSP, SD. LP guarantees the quality and accuracy of the results presented.

## Data sharing

Access to data for authorised researchers is provided within the UCL data safe haven (https://www.ucl.ac.uk/isd/itforslms/services/handling-sens-data) for researchers who have undergone data safe haven and information governance training. Linked CALIBER data (primary care data, Hospital Episode Statistics and Office for National Statistics mortality data) were obtained from the Clinical Practice Research Datalink (www.cprd.com). Access to data is only available once approval has been obtained through the individual constituent entities controlling access to the data. The phenotype algorithms described in this paper are freely available via the CALIBER website at www.caliberresearch.org and the CALIBER data portal is available for consultation online at http://www.caliberresearch.org

## Competing interests

All authors have completed the ICMJE uniform disclosure form at www.icmje.org/coi_disclosure.pdf and declare: no support from any organisation for the submitted work; no financial relationships with any organisations that might have an interest in the submitted work in the previous three years; no other relationships or activities that could appear to have influenced the submitted work; JTT has received research grant funding from Pfizer-BMS (manufacturer of a direct oral anticoagulant) for a completed clinical trial on atrial fibrillation in stroke; AB has been an advisory panel member for Boehringer Ingelheim and Novo Nordisk in the last 3 years;

