## Supplementary Appendix for "Bleeding in cardiac patients prescribed antithrombotic drugs: Electronic health record phenotyping algorithms, incidence, trends and prognosis"

| eMethods | Development of the CALIBER bleeding EHR phenotype algorithm |
| --- | --- |
| eTable 1 | Summary of previous bleeding EHR phenotype algorithms developed using electronic health records |
| eTable 2 | Bleeding terms from Read and ICD-10 |
| eTable 3 | Comparison of results of clinicians review vs. bleeding EHR phenotype for identifying bleeding events |
| eTable 4 | Patient baseline characteristics (time of AF, MI, UA or SA diagnosis) stratified by first CALIBER bleeding event type |
| eTable 5 | Bleeding definitions used in clinical practice |
| eFigure 1 | Overlap of 39804 bleeding records in CPRD (primary care), HES (secondary care) and ONS (death registry) and the number of inferred bleeding cases in patients without a bleeding record in primary or secondary care (n= 128815 patients) |
| eFigure 2 | Five year risk of recurrent bleeding stratified by initial bleeding type: any bleeding or bleeding with further markers of severity (bleeding +). A: Risk of any bleeding; B: Risk of fatal or bleeding with further markers of severity |
| eFigure 3 | Short term mortality with and without indicators of bleeding severity |

**eMethods: Development of the CALIBER bleeding EHR phenotype algorithm**

*1) Reviewing code lists for bleeding and related procedures*

We reviewed bleeding terms in Read and ICD-10 to define bleeding in primary and hospital care and to determine which might be relevant to antithrombotic use. We identified 96 ICD-10 codes and 201 Read codes (**eTable 3**) for bleeding events which we categorised into 18 anatomical sites [Intracranial (intracerebral, subarachnoid, extradural, subdural, unspecified); Gastrointestinal (upper, lower, unspecified); Respiratory (upper, lower, unspecified); Ruptured aortic aneurysm or haemopericardium; Genitourinary; Bleeding disorder; Ocular; Ear; Renal; Unspecified]. We used the field related to bleeding complications in MINAP, which consists of 6 categories pertaining to location and haemoglobin drop. Furthermore we reviewed primary care Read codes and hospital care OPCS codes for procedures including transfusion, haematoma evacuation or aspiration, surgical arrest for bleeding and endoscopy. All codes used to derive the bleeding phenotype are available at the CALIBER portal **[**<https://www.caliberresearch.org/portal>].

*2) Preliminary analysis*

We performed a preliminary analysis of the bleeding records in our study population to determine which data are viable for use in the final algorithm. First we examined the characteristics of the population at entry by whether they had a bleeding or transfusion code in primary or hospital care, or bleeding listed as a cause of death in follow up. We examined the characteristics of bleeding records stratified by anatomical bleeding site. For hospitalised bleeding we calculated the length of hospitalisation and the proportion of records with primary diagnosis position. We investigated the presence of procedures records (transfusion, bleeding surgical arrest, haematoma evacuation and endoscopy) at different time intervals (on the same day, within 7 days and within 30 days) of bleeding records in primary and hospital care. Haemoglobin drop related to a bleeding event was calculated as the peak haemoglobin value within 365 to 7 days prior to bleeding minus the lowest haemoglobin value recorded within 7 days of bleeding for patients with a minimum of 2 haemoglobin values recorded in primary care.

*3) Markers of bleeding severity*

To assess the suitability of markers for bleeding severity we examined short term (30 and 90 days) all-cause and bleeding specific mortality following bleeding with and without the indicators. Guided by current clinical bleeding definitions **(eTable 4)** and availability of data within CALIBER the following severity indicators were considered, 1) anatomical site, 2) presence of a transfusion record within 30 days, 3) presence of surgical interventions within 30 days, 4) haemoglobin drop and 5) bleeding from more than one site on a single date. Furthermore for hospitalised bleeding records we considered primary versus secondary diagnosis position and length of hospitalisation.

The Kaplan-Meier analysis of severity markers and short term outcomes is shown in **eFigure 3**. These were used to inform the definition of severity in the algorithm. For example, transfusion records in primary care were uncommon and only showed modest signs of associated increased short term mortality and therefore only transfusions recorded in hospital care are used to define severity in the algorithm**.**

*4) Inferring bleeding cases*

We attempted to capture potential bleeding cases in patients with no bleeding code in primary or hospital care through the following pathways:

- Surgical procedures (surgical arrest, haematoma evacuation) recorded in primary and hospital care
- Transfusion with
  - Iron deficiency anaemia record in primary care or hospital within 30 days
  - Low haemoglobin (<10g/dL) and endoscopy within 30 days and no cancer, liver disease, renal disease diagnosis 1 year prior
- Low haemoglobin (<10g/dL) with
  - Iron deficiency anaemia record in primary care or hospital care and endoscopy within 30 days and no cancer, liver disease, renal disease diagnosis 1 year prior

*5) Developing the phenotype*

Based on a combination of exploratory analyses of bleeding codes and severity markers and consensus amongst the study team we iteratively developed an algorithm to define bleeding in linked EHR. We grouped bleeding as major and minor both within primary and hospital care, or inferred.

We assessed how well bleeding events are captured amongst the data sources used, allowing up to 30 days between bleeding events in different sources to be considered the same event, using a Venn diagram.

**eTable 1: Summary of previous bleeding EHR phenotypes developed in electronic health records**

| **Author** | **Year** | **Bleeding endpoint(s) evaluated** | **Data source(s)** | **Setting** | **Study Population** | **Coding system**  **(n codes)** | **Supporting EHR data used in case definition** | **Algorithm figure reported** | **Assessment of phenotype accuracy** |
| --- | --- | --- | --- | --- | --- | --- | --- | --- | --- |
| Raiford et al^1^ | 1996 | Upper GI bleeding or perforation | Saskatchewan Hospital Services Plan | Hospital admissions | Patients hospitalised for upper GI bleeding | ICD-9 (30) | No | No | Site specific codes  PPV: 91%  Nonspecific codes  PPV: 68% |
| De Abajo et al^2^ | 1999 | Upper GI bleeding | GPRD  (UK) | Primary care | Patients with a record for acute upper GI bleeding | Read (codes not stated) | No | No | PPV: 95/96 |
| Arnason et al^3^ | 2006 | From 8 anatomical sites:  Any bleeding  Major bleeding | A university hospital, Ottawa | Hospital admissions | Patients with a record for thromboembolism or bleeding | ICD-9 (81) | No  (information from patient charts were used to classify severity) | No | Definite bleeding  PPV: 91%; NPV: 91%  Major bleeding  PPV: 87%; NPV: 92% |
| Wahl et al^4^ | 2010 | Severe upper GI bleeding | HealthCore Integrated Research Database (USA) | Hospital admissions | Patients with a record for upper GI bleeding | ICD-9 (original:75; refined:33) | Procedure codes | No | PPV:  original: 56.5%  refined: 87.8% |
| Cunningham et al^5^ | 2011 | Serious bleeding related to oral anticoagulation from >4 anatomical sites | Tennessee Medicaid program | Hospital admissions | Medicaid enrolees >30 years old | ICD-9 (39) | No  (information from patient charts were used to classify severity) | No | PPV assessed for individual codes ranged from 71.4% to 100% (>5 charts) |
| Crooks et al^6^ | 2012 | Upper GI bleeding | CPRD  HES  ONS  (UK) | Primary care  Hospital admissions  Death registry | Patients with a record for acute upper GI bleeding | Read (46)  ICD-10 (22) | Causes, symptoms, endoscopy, death, transfusion, procedures, alcohol, anaemia, coagulation, collapse, other | Yes | None |
| Valkhoff et al^7^ | 2014 | Upper GI bleeding | IPCI (Netherlands)  HSD (Italy)  ARS (Italy)  Aarhus (Denmark) | Primary care  Hospital admissions | Patients with a record for upper GI bleeding | IPCI (4)  ICD-9 (26)  ICD-10 (16) | No | No | IPCI - PPV: 21%  HSD - PPV: 78%  ARS - PPV: 72%  Aarhus - PPV: 77% |
| Friberg et al^8^ | 2016 | In 4 categories of anatomical site:  Fatal  Non-fatal major  Hospitalised  Minor | Swedish Patient register | Hospital admissions  Hospital outpatients  Death registry | Atrial fibrillation patients | ICD-10 (38) | Anatomical site  (intracranial)  Transfusion  Hospitalisation  Diagnosis position | No | Fatal  PPV: 88.1%; NPV: 99.7%  Non-fatal major  PPV: 90.6%; NPV: 91.5%  Hospitalised  PPV: 65.1%; NPV: 97.5%  Minor  PPV: 84.2%; NPV: 98.9% |
| **CALIBER – present study** | 2017 | In 18 categories of anatomical site:  Fatal  Hospitalised with markers of severity  Hospitalised  Primary care with markers of severity  Primary care  Inferred | CPRD  HES  ONS | Primary care  Hospital admissions  Death registry | Coronary disease and atrial fibrillation patients | Read (201)  ICD-10(96) | Yes: Transfusion, anatomical site; procedures; | Yes | Analysed prognosis following bleeding |

Note: GI= gastrointestinal; PPV= positive predictive value; NPV= negative predictive value

**eTable 2: Bleeding Read and ICD-10 codes**

| **Anatomical Site** | **ICD-10 term** | **Read codes** |
| --- | --- | --- |
| Intracranial (Intracerebral) | I61; I610; I611; I612; I613; I614; I615; I616; I618; I619; | G61z.00; G61..00; G614.00; G613.00; G61X100; G61X000; G600.00; G616.00; G617.00; G61X.00; G610.00; G611.00; G612.00; Gyu6200; G618.00; G615.00; Gyu6F00 |
| Intracranial (Subarachnoid) | I60; I600; I601; I602; I603; I604; I605; I606; I607; I608; I609; I62 | G60..00; G604.00; G602.00; G60z.00; G605.00; G603.00; G601.00; G606.00; Gyu6100 |
| Intracranial (Subdural) | I620 | G621.00; G623.00 |
| Intracranial (extradural) | I621; S064 | G620.00; S626.00 |
| Intracranial (unspecified) | I629 | G62z.00; G62..00 |
| Aortic aneurysm or haemopericardium | I230; I312; I711; I713; I715; I718 | G715000; G713.11; G715.00; G530.00; G711.11; G713.00; G360.00; G711.00; G723500; G713000 |
| Upper gastrointestinal | I850; I983; K250; K252; K254; K256;K260; K262; K264; K266; K270; K272; K274; K276; K280; K282; K284; K286; K290; K661; K920 | J110111; J120100; J150000; J110100; J130100; J130300; J120300; J110300; J140100; J680.00; J680.11; J12y100; J68z200; J68z000; J10y000; 1994.11; J121111; 7619100; G850.00; J12yy00; J111111; 4A5..11; 4A23.11; 1994; 1995; J121100; J131100; 4A23.00; J11y100; 4A5..00; J14y100; 4A51.00; J111100; J13y100; 4A5Z.00; J121300; J111300; J12y300; J11yy00; J13y300; G852000 |
| Lower gastrointestinal | K625; K921 | J681.00; J573011; J510900; J68z100; J681.11; SE23111; 19E6.00; 19E6.11; J573012; J573000; SE22300; 196C.00; 196B.00; 479..11; 4762.11; 19E4.12; J573.00; J681.13; J681.12; J573100; 4737.11; 4762; S740100; J573z00 |
| Unspecified gastrointestinal | K922 | J68z.11; J68..00; J68zz00; J68z.00 |
| Genitourinary | N02; N026; N028; N029; N421; N501; N836; N837; N857; N897; N898; N908; N921; N925; N926; N93; N938; N939; N950; R31; R31X | K197.00; K286v00; K5E..00; 1A45.00; K197400; K197300; K19y411; 1584; K197000; K19y400; K59yx00; K56y100; K286w00; K575.00; K197100; K5E1.00; K286100; K5E2.00; K59yy00; K5E0.00; K5Ez.00; K566.00; K55y300; K16y200; K275100; K221100; K286400; K537.00; K167.00; K275200; Kyu9D00 |
| Upper respiratory | R040; R041 | R047.00; R047.11; 1C62.00; 2D25.00; R048.00; 2DE7.00 |
| Lower respiratory | J942; R042 | R063.00; 172..00; R063100; R063000; R063z00 |
| Unspecified respiratory | R048; R049 | - |
| Ocular | H313; H356; H431; H450 | F4K7.00; F4K2800; F404500; F42y500; 2BB8.00; 2BB5.00; F42y.11; F436000; F4Ey000; F436100; F42y400; FyuH400; F436.00; F4H4100; F42y100; F436z00; F424300; F437200; F42y300 |
| Ear | H922 | F503100; F501G00 |
| Renal | - | S760100; K13y800; S760111; K138300; K138100; S761100; C154200 |
| Bleeding disorders | D683; D69; D698; D699 | D31..00; D31z.00; D31X.00; D31yz00; D31y.00; Dyu3300 |
| Unspecified | R233; R58; R58X; T81 | SE...11; SE33011; SE4z.11; SE45.11; SK02.12; SK02.00; SE4z.12; S750100; 2F65.00; SK02.11; S751100; Ryu7300 |

**eTable 3: Comparison of results of clinicians review vs. bleeding EHR phenotype for identifying bleeding events**

| **Bleeding phenotype algorithm** | **Clinician review** | |  |
| --- | --- | --- | --- |
|  | Bleeding | No bleeding | Total |
| Bleeding | 15 | 2 | 17 |
| No bleeding | 16 | 250 | 266 |
| Total | 31 | 252 | 283 |

**eTable 4: Patient baseline characteristics (time of Atrial fibrillation, Myocardial infarction, Unstable angina or Stable angina diagnosis) stratified by first CALIBER bleeding event type**

|  |  | **I** | **II** | | | **III** |  |
| --- | --- | --- | --- | --- | --- | --- | --- |
|  | **Fatal (n=1575)** | **Hospitalised +MS**  **(n=2654)** | **Primary care +MS (n=1645)** | **Hospitalised (n=8766)** | **Inferred**  **(n=1144)** | **Primary care (n=12619)** | **No bleeding (n=100412)** |
| **Demographics and behaviours**  **at cohort entry** |  |  |  |  |  |  |  |
| Age (years), mean (SD) | 77.4 (9.76) | 75.6 (10.98) | 73.6 (10.81) | 71.5 (11.86) | 74.1 (11.48) | 70.1 (11.88) | 71.4 (13.31) |
| Women, n (%) | 690 (43.8) | 1224 (46.1) | 680 (41.3) | 3977 (45.4) | 549 (48.0) | 5124 (40.6) | 46762 (46.6) |
| Highest quartile of deprivation (most deprived) | 341 (21.7) | 526 (19.9) | 292 (17.8) | 1922 (22.0) | 239 (20.9) | 2322 (18.4) | 20037 (20.0) |
| *% missing* | 0.2 | 0.3 | 0.1 | 0.3 | 0.3 | 0.2 | 0.3 |
| Smoking status, n (%) |  |  |  |  |  |  |  |
| Non-Smoker | 631 (52.6) | 1149 (56.9) | 718 (57.3) | 3751 (54) | 484 (54.4) | 5527 (54.9) | 41145 (51.1) |
| Current smoker | 160 (13.3) | 183 (9.1) | 97 (7.7) | 785 (11.3) | 104 (11.7) | 1006 (10.0) | 10920 (13.6) |
| Ex-smoker | 409 (34.1) | 688 (34.1) | 439 (35.0) | 2413 (34.7) | 301 (33.9) | 3535 (35.1) | 28478 (35.4) |
| *% missing* | 23.8 | 23.9 | 23.8 | 20.7 | 22.3 | 20.2 | 19.8 |
| History of alcohol abuse, n (%) | 125 (7.9) | 265 (10.0) | 159 (9.7) | 868 (9.9) | 115 (10.1) | 1203 (9.5) | 9689 (9.6) |
| **Medical history**  ***(Any record ever prior to entry)*** |  |  |  |  |  |  |  |
| Myocardial infarction, n (%) | 290 (18.4) | 486 (18.3) | 296 (18.0) | 1508 (17.2) | 190 (16.6) | 2098 (16.6) | 21778 (21.7) |
| Atrial fibrillation, n (%) | 411 (26.1) | 677 (25.5) | 303 (18.4) | 1685 (19.2) | 295 (25.8) | 1870 (14.8) | 21820 (21.7) |
| Stable angina, n (%) | 820 (52.1) | 1372 (51.7) | 959 (58.3) | 5115 (58.4) | 613 (53.6) | 7930 (62.8) | 52960 (52.7) |
| Unstable angina, n (%) | 129 (8.2) | 251 (9.5) | 157 (9.5) | 900 (10.3) | 114 (10.0) | 1271 (10.1) | 9073 (9.0) |
| Diabetes, n (%) |  |  |  |  |  |  |  |
| Type 1 | 10 (0.6) | 31 (1.2) | 21 (1.3) | 95 (1.1) | 11 (1.0) | 112 (0.9) | 858 (0.9) |
| Type 2 | 164 (10.4) | 367 (13.8) | 208 (12.6) | 1124 (12.8) | 180 (15.7) | 1406 (11.1) | 12118 (12.1) |
| Unspecified type | 39 (2.5) | 58 (2.2) | 30 (1.8) | 202 (2.3) | 28 (2.4) | 198 (1.6) | 1951 (1.9) |
| Stroke (ischaemic or unspecified), n (%) | 133 (8.4) | 251 (9.5) | 97 (5.9) | 527 (6.0) | 80 (7.0) | 621 (4.9) | 5915 (5.9) |
| Peripheral arterial disease, n (%) | 205 (13.0) | 312 (11.8) | 174 (10.6) | 850 (9.7) | 136 (11.9) | 1068 (8.5) | 8742 (8.7) |
| Renal disease, n (%) | 143 (9.1) | 235 (8.9) | 144 (8.8) | 659 (7.5) | 91 (8.0) | 658 (5.2) | 7416 (7.4) |
| Cancer, n (%) | 233 (14.8) | 477 (18.0) | 366 (22.2) | 1354 (15.4) | 194 (17.0) | 1777 (14.1) | 14040 (14.0) |
| Peptic ulcer, n (%) | 142 (9.0) | 244 (9.2) | 173 (10.5) | 792 (9.0) | 78 (6.8) | 1023 (8.1) | 6902 (6.9) |
| Bleeding diatheses or coagulation disorders, n (%) | 25 (1.6) | 38 (1.4) | 31 (1.9) | 103 (1.2) | 15 (1.3) | 109 (0.9) | 777 (0.8) |
| Chronic anaemia, n (%) | 260 (16.5) | 491 (18.5) | 453 (27.5) | 1305 (14.9) | 372 (32.5) | 1303 (10.3) | 12929 (12.9) |
| **Biomarkers**  ***(Nearest record to entry within 1 year prior)*** |  |  |  |  |  |  |  |
| Systolic blood pressure (mmHg), mean (SD) | 143 (21.5) | 144 (21.4) | 144 (22.2) | 142 (21.1) | 142 (21.6) | 143 (20.8) | 141 (20.9) |
| *% missing* | 26.0 | 27.9 | 24.9 | 26.2 | 26.1 | 23.7 | 26.1 |
| Haemoglobin (g/dL), mean(SD) | 13.1 (1.90) | 12.9 (1.96) | 12.5 (2.19) | 13.4 (1.81) | 11.7 (2.74) | 13.7 (1.60) | 13.4 (1.80) |
| *% missing* | 62.6 | 61.6 | 59.5 | 61.7 | 57.5 | 62.2 | 59.6 |
| Creatinine (mol/l), median (IQR) | 105 (87, 129) | 100 (84, 122) | 101 (86, 123) | 96 (82, 114) | 101 (85, 123) | 95 (82, 111) | 94 (81, 112) |
| Min, Max | 23.00, 733 | 4.00, 1290 | 8.60, 906 | 1.00, 1036 | 45.00, 739 | 2.60, 919 | 0.11, 1625 |
| *% missing* | 53.5 | 55.1 | 53.1 | 54.1 | 50.6 | 54.2 | 50.9 |
| Body mass index, mean (SD) | 27.0 (5.57) | 27.4 (5.49) | 28.2 (6.07) | 28.3 (5.66) | 27.3 (4.90) | 28.5 (5.44) | 28.1 (5.76) |
| Underweight | 15 (3.1) | 21 (2.6) | 11 (1.9) | 51 (1.7) | 8 (2.0) | 60 (1.4) | 893 (2.6) |
| Normal | 161 (33.8) | 271 (33.2) | 151 (26.5) | 792 (27.1) | 122 (31.2) | 1084 (24.6) | 9416 (27.9) |
| Overweight | 192 (40.3) | 308 (37.7) | 237 (41.7) | 1179 (40.4) | 166 (42.5) | 1824 (41.4) | 12775 (37.8) |
| Obese | 109 (22.9) | 216 (26.5) | 170 (29.9) | 897 (30.7) | 95 (24.3) | 1433 (32.6) | 10675 (31.6) |
| *% missing* | 69.7 | 69.3 | 65.4 | 66.7 | 65.8 | 65.1 | 66.4 |
| **Prescribed antithrombotic therapies and duration**  **between cardiac disease diagnosis and 1^st^ bleeding event (median, IQR)** |  |  |  |  |  |  |  |
| No antithrombotic therapy, n (%) | 436 (27.7) | 591 (22.3) | 402 (24.4) | 1554 (17.7) | 453 (39.6) | 2012 (15.9) | 22983 (22.9) |
| Aspirin monotherapy, n (%) | 852 (54.1) | 1547 (58.3) | 961 (58.4) | 5473 (62.4) | 514 (44.9) | 8344 (66.1) | 64564 (64.3) |
| Duration (days) | 619 (180, 1317) | 538 (179, 1247) | 459 (163, 1098) | 524 (170, 1184) | 427 (127, 1018) | 557 (185, 1228) | 820 (288, 1768) |
| Clopidogrel monotherapy, n (%) | 128 (8.1) | 209 (7.9) | 139 (8.4) | 876 (10.0) | 76 (6.6) | 1264 (10.0) | 11031 (11.0) |
| Duration (days) | 131 (40.5, 550) | 128 (38.0, 422) | 174 (46.0, 558) | 117 (35.0, 448) | 110 (58.8, 447) | 121 (40.0, 440) | 146 (43.0, 567) |
| Dual antiplatelet therapy, n (%) | 166 (10.5) | 339 (12.8) | 188 (11.4) | 1330 (15.2) | 115 (10.1) | 1843 (14.6) | 18242 (18.2) |
| Duration (days) | 216 (90.0, 476) | 174 (77.5, 402) | 164 (80.2, 398) | 197 (90.0, 423) | 148 (77.0, 376) | 180 (90.0, 402) | 322 (118.0, 486) |
| VKA monotherapy, n (%) | 261 (16.6) | 479 (18.0) | 239 (14.5) | 1439 (16.4) | 133 (11.6) | 1892 (15.0) | 11512 (11.5) |
| Duration (days) | 274 (99, 892) | 411 (108, 916) | 280 (99, 656) | 313 (105, 806) | 267 (90, 782) | 292 (97, 775) | 387 (132, 1036) |
| VKA + 1 antiplatelet, n (%) | 139 (8.8) | 205 (7.7) | 121 (7.4) | 740 (8.4) | 59 (5.2) | 1038 (8.2) | 6656 (6.6) |
| Duration (days) | 76 (41.0, 171) | 90 (49.0, 218) | 90 (57.0, 180) | 90 (48.0, 232) | 104 (61.5, 434) | 90 (52.0, 217) | 90 (52.0, 200) |
| VKA + 2 antiplatelets, n (%) | 20 (1.3) | 26 (1) | 15 (0.9) | 77 (0.9) | 3 (0.3) | 104 (0.8) | 874 (0.9) |
| Duration (days) | 70.0 (34.8, 91.0) | 41.5 (19.2, 68.8) | 44.0 (16.0, 79.5) | 59.0 (38.0, 90.0) | 69.0 (44.5, 79.5) | 57.5 (35.0, 90.0) | 64.0 (40.0, 90.0) |

Note: MS= markers of severity; SD= standard deviation; IQR= interquartile range; VKA= vitamin K antagonist

**eTable 5: Bleeding definitions used in clinical trials and observational studies and factors used to classify severity**

| Factor | CALIBER – present study | Stable post-MI risk prediction^9^ | Bleeding Academic Research Consortium (BARC)^10^ | International Society on Thrombosis and Haemostasis (ISTH)^11^ | Thrombosis In Myocardial Infarction (TIMI)^12^ |
| --- | --- | --- | --- | --- | --- |
| Fatal | ● | ● | ● | ● | ● |
| Anatomic location | ●Intracranial; Ruptured aortic aneurysm; Haemopericardium | ●Intracranial | ●Intracranial; Intraocular | ● Intracranial; Intraspinal; Intraocular; Retroperitoneal; Intraarticular; Pericardial; Intramuscular w. compartment syndrome | ● Intracranial |
| Haemoglobin drop | ○ | ○ | ● 3 - <5g/dL; ≥5g/dL | ● ≥2 g/dL | ● 3 - <5g/dL; ≥5g/dL |
| Hospitalisation | ● | ● | ● | ● | ○ |
| Mode of hospital admission | ● | ○ | ○ | ○ | ○ |
| Length of hospitalisation | ● >14 days | ● > 14 days | ○ | ○ | ○ |
| Blood transfusion | ● | ● | ● | ● | ● |
| Number of units transfused | ○ | ○ | ● | ● | ○ |
| Medical/surgical consultation | ● | ○ | ● | ● | ● |
| Medical or surgical intervention | ● | ○ | ● | ● | ● |
| Multiple bleeding codes | ● | ○ | ○ | ○ | ○ |
| Haemodynamic compromise | ○ | ○ | ○ | ● | ○ |
| Change in antithrombotic therapy | ○ | ○ | ○ | ● | ● |

**eFigure 1: Overlap of 39,804 bleeding recorded in CPRD (primary care), HES (hospital admissions) and ONS (death registry) and the number of inferred bleeding cases in patients without a bleeding record in primary or hospital care (n= 128,815 patients)**

67

4689

445

60

**HES**

**CPRD**

**ONS**

Bleeding recorded

Bleeding Inferred

**Note: Numbers of inferred (possible) bleeding events according to source of information: 477 surgical arrest or haematoma evacuation procedures in 451 patients; in 514 patients 689 cases of a transfusion code in OPCS accompanied by an iron deficiency anaemia diagnosis in HES or CPRD within 30 days; in 62 patients 77 cases of a transfusion code in OPCS accompanied by a haemoglobin value of <10g/dL in CPRD within 30 days, an endoscopic examination within 30 days and no history of cancer, liver or renal disease in the year prior to transfusion; and in 182 patients 249 cases of haemoglobin <10g/dL in CPRD, an endoscopic examination within 30 days and no history of cancer, liver or renal disease in the year prior to the haemoglobin record. That is, overall 1,492 potential bleeding events identified in 1144/101,566 (1.1%) patients with no bleeding record in HES or CPRD.
eFigure 2: Five year risk of recurrent bleeding stratified by initial bleeding type: any bleeding or bleeding with further markers of severity (bleeding +). A: Risk of any bleeding; B: Risk of fatal or bleeding with further markers of severity**

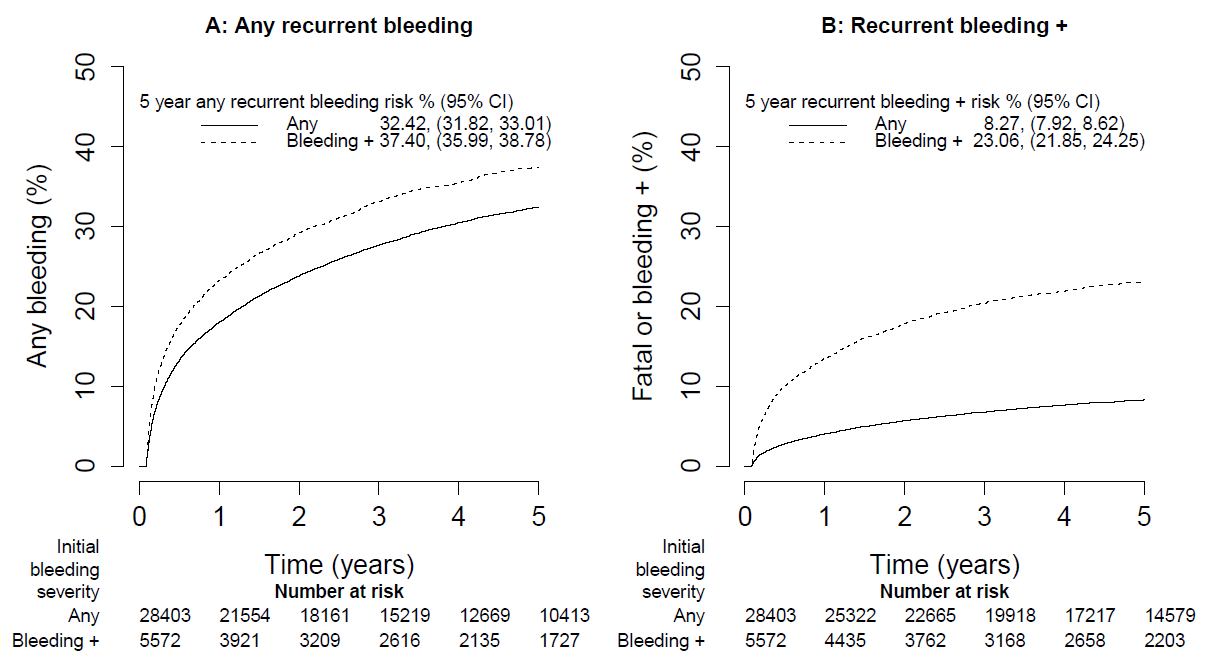

**Note:** ‘Any bleeding’ includes hospitalised, hospitalised +, primary care, primary care + and inferred bleeding. ‘Bleeding +’ includes hospitalised + or primary care + bleeding

**eFigure 3: Short term mortality with and without indicators of bleeding severity**

| Bleeding location | 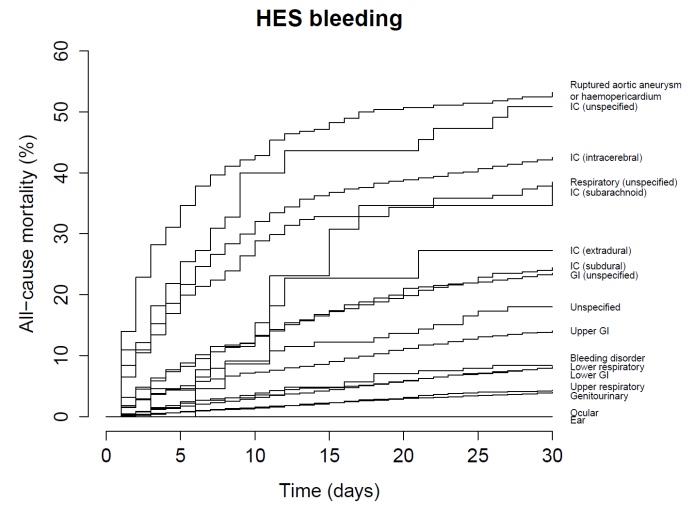 | 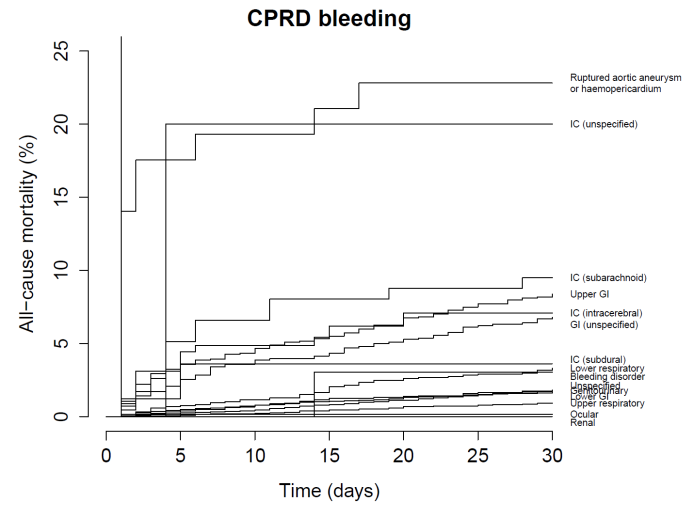 |
| --- | --- | --- |
| Hospitalisation diagnosis position | 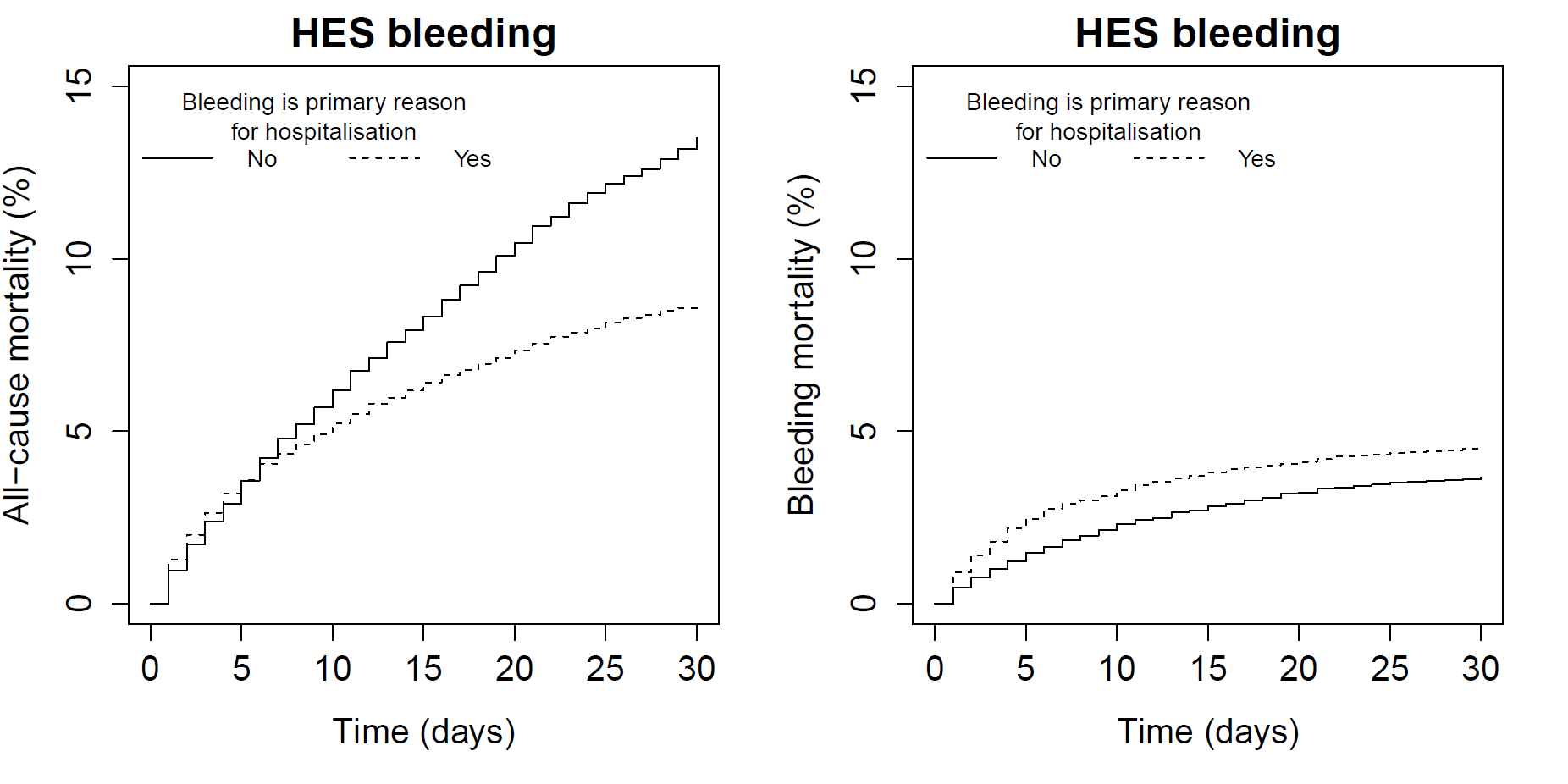 | 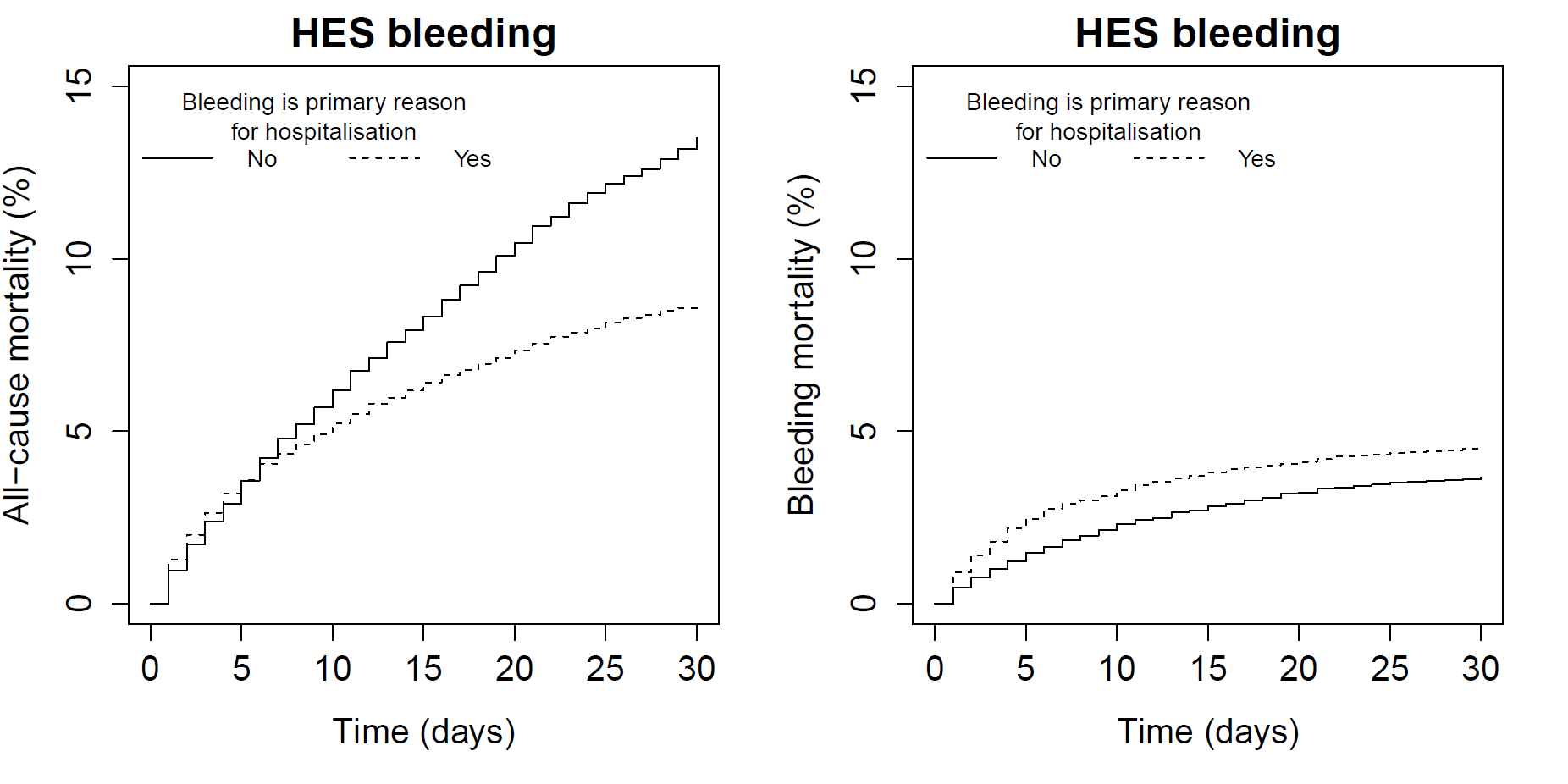 |
| Transfusion | 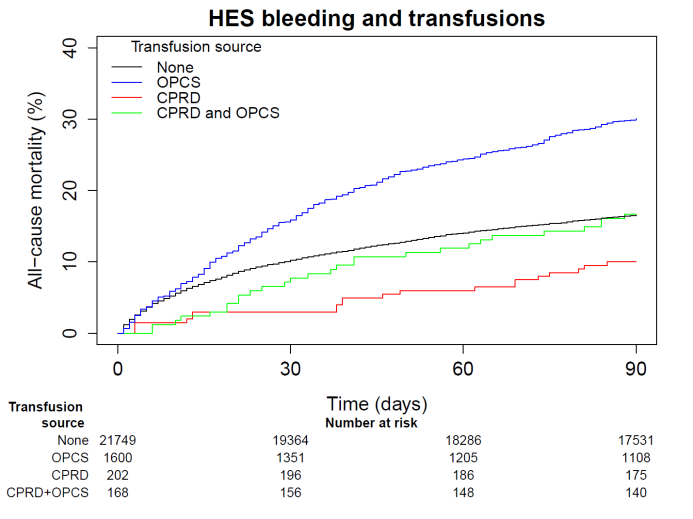 | 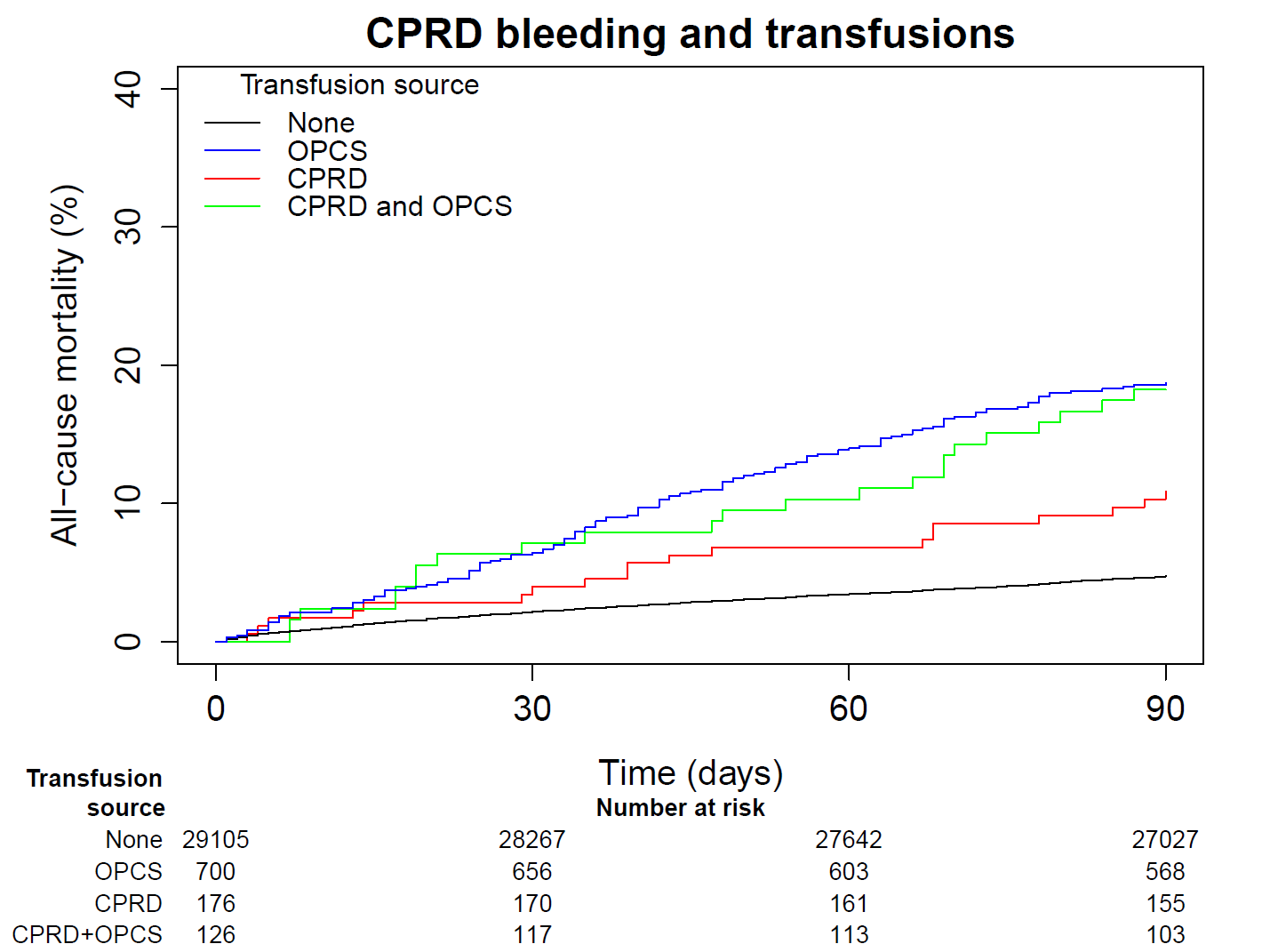 |
| Haemoglobin drop | 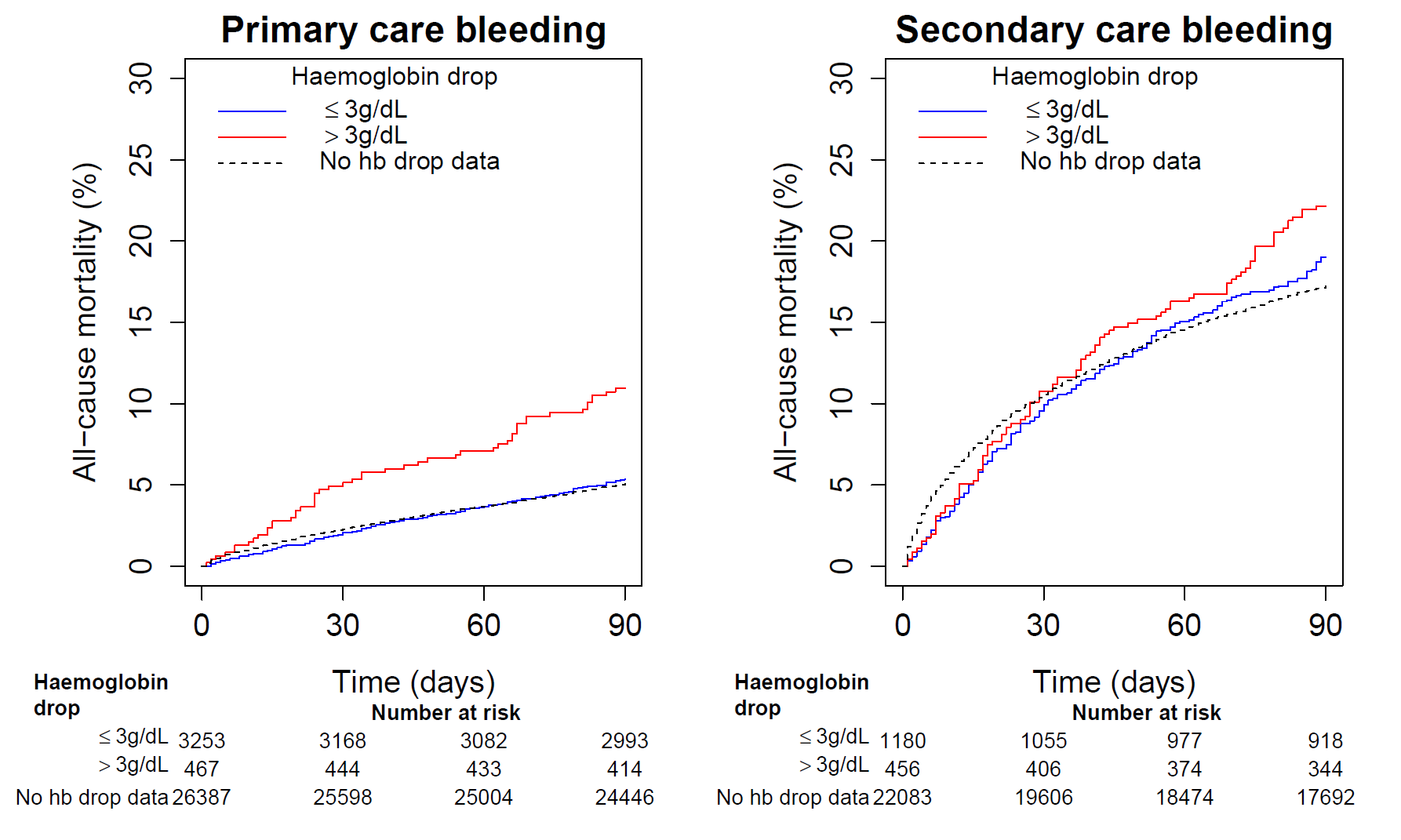 | 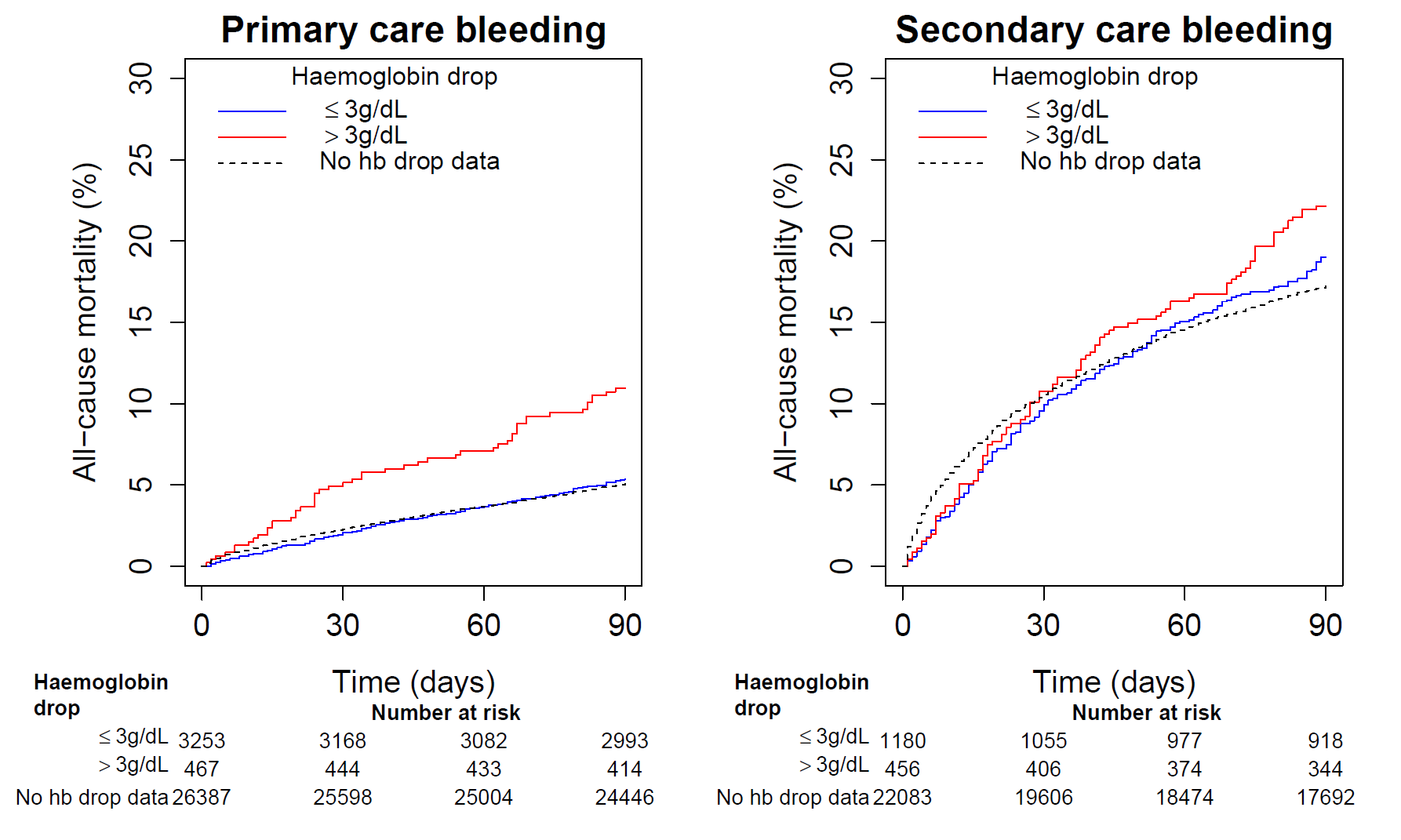 |
| Number of bleeding codes | 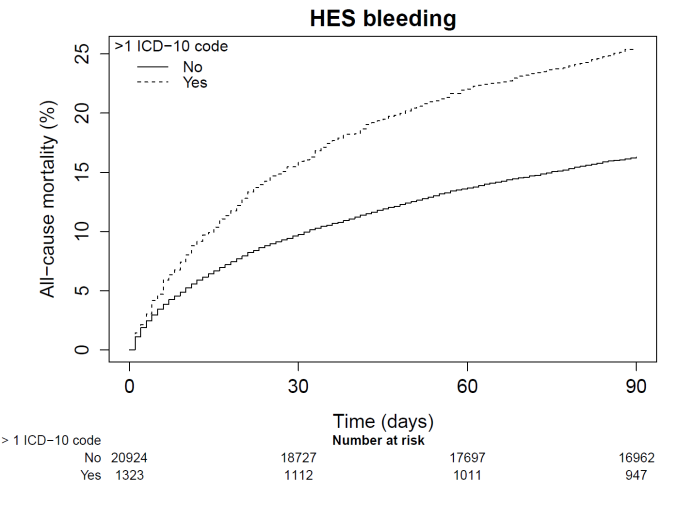 | 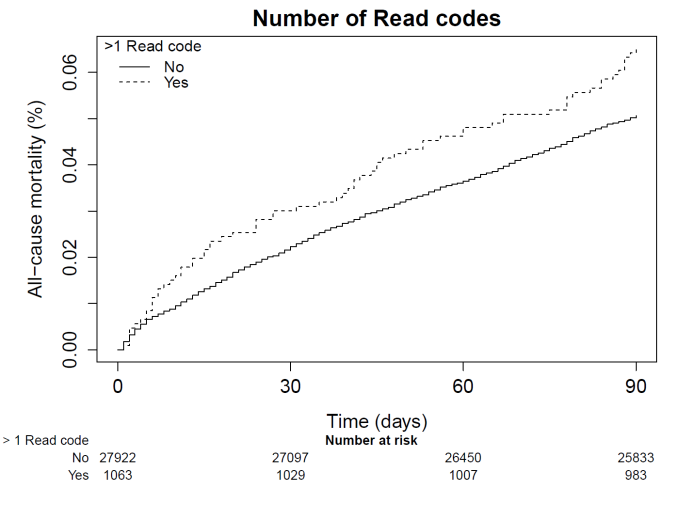 |
| Endoscopy | 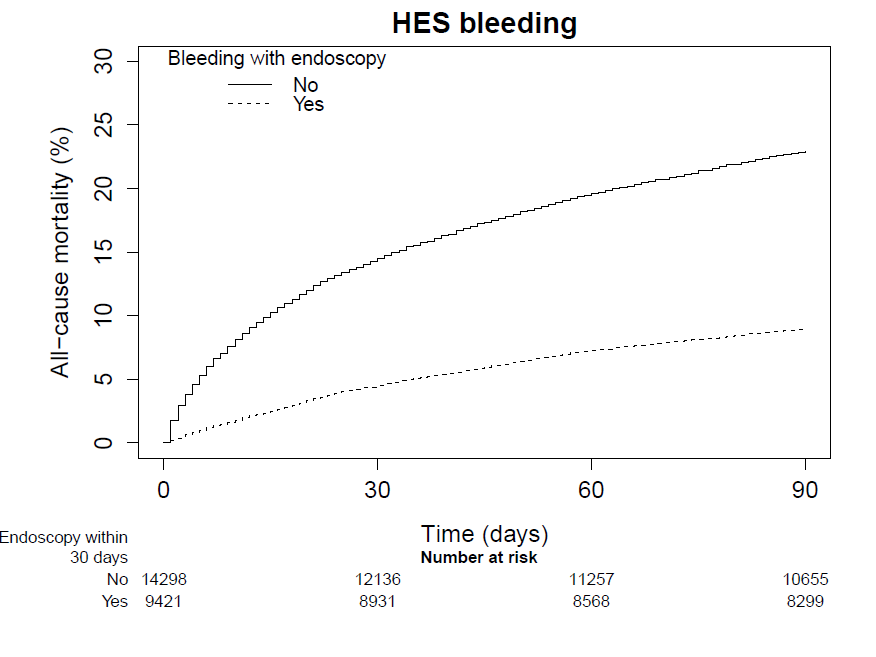 |  |
| Bleeding intervention procedures | 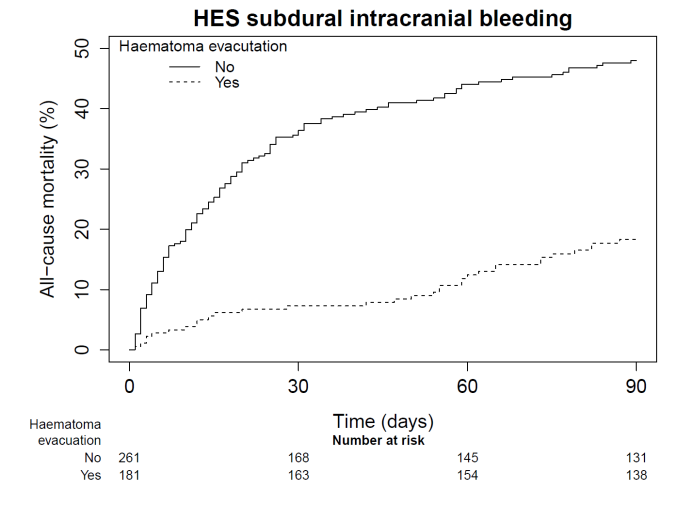 | 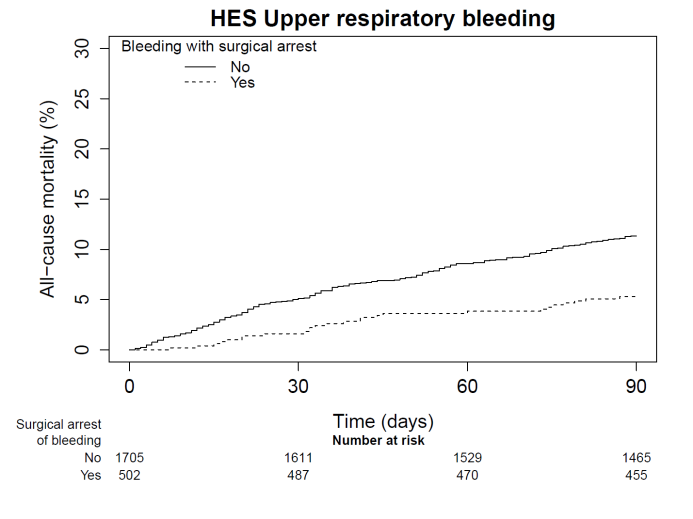 |
